# Synthetic inhibition of the SUMO pathway by targeting the SAE1 component via TAK-981 compound impairs tumor growth and chemosensitizers embryonal and alveolar rhabdomyosarcoma cell lines

**DOI:** 10.1101/2025.01.15.633172

**Authors:** Silvia Codenotti, Volker M. Lauschke, Emma V. Casella, Daniel C. Andersson, Alessandro Fanzani, Stefano Gastaldello

## Abstract

**Background:** Rhabdomyosarcoma (RMS) is a highly aggressive pediatric soft tissue sarcoma with limited therapeutic options, particularly for cases resistant to conventional treatments. The SUMOylation pathway, which plays a key role in regulating cell cycle, apoptosis, and transcription, has emerged as a potential therapeutic target in RMS. Elevated levels of SUMO1 and SUMO2/3 conjugates in RMS cell lines, compared to normal human skeletal muscle cells, underscore the association between hyper-SUMOylation and aggressive cancer phenotypes. Understanding these molecular underpinnings is critical for the development of innovative and effective treatments.

**Methods:** The investigation encompassed transcriptomic and protein analyses to profile SUMOylation pathway components across alveolar and embryonal RMS subtypes, aiming to identify heterogeneity that could guide personalized therapy approaches. TAK-981’s interactions with chemotherapeutic agents, were evaluated for synergistic effects. Additionally, its impact on radiosensitivity and key signaling pathways, such as AKT and ERK phosphorylation, was assessed to elucidate its mechanism of action.

**Results:** Transcriptomic and proteomic analyses revealed distinct expression profiles of SUMOylation pathway components across RMS subtypes, highlighting heterogeneity that could guide personalized therapeutic strategies. Notably, SAE1 protein was overexpressed in RMS tissues and cells, positioning it as a potential biomarker for this cancer. Its activity was effectively counteracted by TAK-981, a SUMO inhibitor that demonstrated significant therapeutic potential by suppressing RMS cell proliferation and migration and enhancing the cytotoxic effects of chemotherapeutic agents, actinomycin D and doxorubicin. However, TAK-981 did not increase radiosensitivity, suggesting its selective action through chemical inhibition mechanisms. Mechanistically, TAK-981 reduced phosphorylation of key signaling proteins, including AKT and ERK, critical for RMS cell survival.

**Conclusion:** The findings of this study establish TAK-981 as a promising therapeutic agent for RMS. The results also provide foundational insights into the role of SUMOylation associated to the new biomarker SAE1 in RMS and its subtypes, paving the way for the development of personalized treatment strategies that leverage SUMO pathway inhibition.

## 1. Introduction

Rhabdomyosarcoma (RMS) is a rare and aggressive form of cancer that originates in the soft tissues, specifically in skeletal muscle cells. This malignancy primarily affects children and adolescents, though it can also occur in adults. RMS is thought to arise from embryonic muscle cells that fail to differentiate into mature muscle cells, leading to the formation of cancerous tissues (1). Based on the presence or absence of the chromosomal translocations t(2;13) or t(1;13) (2) (3), RMS is classified molecularly into fusion-positive RMS (FP-RMS) or fusion-negative RMS (FN-RMS), respectively, giving rise to chimeric factors that play a large role in RMS malignancy (4) (5). Among the six RMS histotypes, the alveolar and embryonal are the most common. The alveolar subtype with an incidence of 32% (6) tends to occur in older children and is generally more aggressive whereas the embryonal subtype is twice as common (6), often found in younger children and associated with a more favorable prognosis. The symptoms of RMS depend on the location of the tumor but may include swelling, pain, or a lump in the affected area. Common sites for these tumors include the head and neck, genitourinary tract, and extremities (7) (8). Treatments strategies for RMS typically involve a multidisciplinary approach, combining surgery, chemotherapy, and radiation therapy (9). Despite advancements in treatment, RMS poses significant challenges, and outcomes can vary.

RMS prognosis varies depending on tumor histotype and status. Patients with low-risk localized disease treated with a multimodal approach have a 5-year survival greater than 70%. The overall survival rate for patients presenting high-risk metastatic or recurrent disease remains lower than 30% (10) (11).

The SUMO network is a crucial cellular mechanism involved in post-translational modification. SUMOylation and de-SUMOylation are reactions that involve the attachment and removal of small ubiquitin-like modifiers (SUMO1-2-3-4-5) to proteins, modifying their function, localization, or interaction with other molecules (12) (13) (14). This reaction is dynamic and reversible, playing a significant role in various cellular activities, including gene expression, DNA repair, and cell cycle regulation. Furthermore, the SUMO network has emerged as an appealing target for pharmacological modulation. SUMOylation begins with the activation of SUMO proteins by an enzyme called E1 (SUMO-activating enzyme). The activated SUMO is then transferred to an E2 enzyme (SUMO-conjugating enzyme). Finally, with the help of E3 enzyme (SUMO ligases-PIAS), SUMOs are conjugated to a lysine residue on the target protein via an iso-peptide bond (15) (16). In contrast, deSUMOylation is primarily carried out by SUMO-specific proteases (SENPs) of the sentrin family, which in humans includes six members (SENP1-3 and SENP5-7) that differ in their tissue-specificity and subcellular localization (17). SENPs cleave the isopeptide bond between SUMO and the target protein, releasing the SUMO molecule from its protein scaffold (18). Together, SUMOylation and deSUMOylation create a dynamic and reversible regulatory system in cells, which is essential for cellular homeostasis, response to stress, and adaptation to changing environmental conditions (19).

Aberrations in the SUMO network have been implicated in cancer development and progression (20) (21). Specifically, the SUMO system is known to modulate the activity of several proteins involved in cancer related pathways, such as those regulating cell cycle progression, apoptosis and DNA repair. In certain cancers such as breast (22), colorectal (23), prostate (24), lung (25), pancreatic (26), the overexpression of specific SUMO enzymes or changes in the SUMOylation status of proteins may promote tumorigenesis (27). Conversely, reduced SUMOylation can compromise the activity of certain tumor suppressor proteins like, BRCA1, PML (Promyelocytic Leukemia Protein) (28). Understanding these intricacies of the SUMO network and its involvement in cancer biology has prompted research into the therapies that target SUMOylation dynamics (29) (30).

There has been interest in developing small molecules that can modulate the SUMO pathway such as Spectomycin B1 (31), Ginkgolic Acid (32), Anacardic Acid (33). N106 (34) and TAK-981 (35) are compounds that specifically enhance or inhibit SUMOylation. In this study, we demonstrated that TAK-981 a mechanism-based inhibitor of SAE which forms a SUMO–TAK-981 adduct as the inhibitory species within the enzyme catalytic site, reduces the SUMOylation in alveolar and embryonal RMS cells in vitro in a dose dependent manner, causing a reduction in cell proliferation and migration. Our data provide the first evidence that interfering of SUMOylation represents a potential targeted therapy to treat patients with SUMOylation gain-of-function RMS. These results thus incentivize further translational investigations of SUMO inhibition in RMS to eventually improve outcomes and minimize side effects.

## 2. Materials and Methods

### 2.1. Chemicals

N-ethylmaleimide (NEM, E1271), sodium chloride (NaCl, Merck 6404), dithiothreitol (DTT, D5545), iodoacetamide (I1149), IGEPAL CA-630 (NP40, I3021), ethylenediaminetetraacetic acid disodium salt dehydrate (EDTA, E4884), sodium deoxycholate monohydrate (DOC, D5670), triton X-100 (T9284), glycine (G-7126), albumin form bovine serum (BSA, A7906), sodium dodecyl sulfate (SDS, L3771), tween-20 (P9416), Trizma base (T93349), actinomycin D (A-9415), and all oligonucleotides for quantitative real-time PCR (Table 1) were purchased from Sigma-Aldrich (St. Louis, MO). Doxorubicin (5927S) was purchased from Cell Signaling Technologies (MA, USA). Complete^TM^ protease inhibitors cocktail tablets (protease inhibitors) and phosphatase inhibitor cocktail were purchased from Roche Diagnostic (Mannheim, Germany).

**Table 1.**
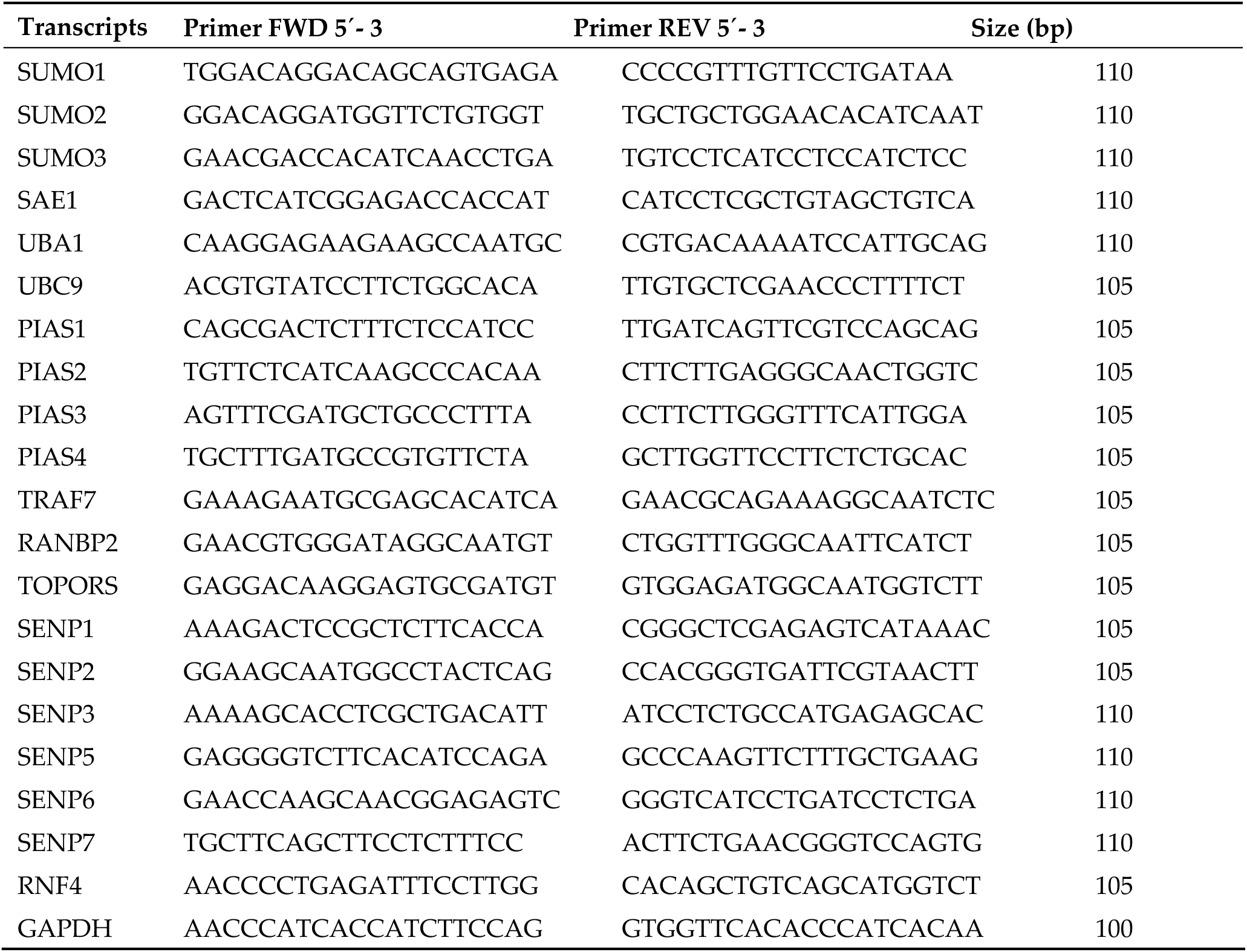

### 2.2. Antibodies

The antibodies used for western blot were: rabbit anti-SUMO-1 (Y299, ab32058, 1:4000), rabbit anti-SUMO-2/3 (EPR4602, ab109005, 1:8000), goat anti-UBE2I/UBC9 (ab21193, 1:3000), rabbit anti-SAE1 (ab185552, 1:2000), rabbit anti-SAE2 (ab185955, 1:5000), rabbit anti-PIAS1 (ab109388, 1:2000), rabbit anti-SENP1 (ab236094, 1:1000), rabbit anti-SENP3 (ab124790, 1:1000), rabbit anti-SENP5 (ab58420, 1:1000), and polyclonal donkey horseradish peroxidase (HRP)-conjugated anti-goat/sheep IgG (ab6885, 1:5000) from AbCam (Cambridge, UK). Rabbit anti-PIAS4 (Pa5-20954, 1:3000) from Thermo Fisher Scientific (IL, USA). Rabbit anti-PIAS3 (SC 14017, 1:2000), mouse anti-phospho-Erk1/2 (sc-7383, 1:500), and mouse anti-Erk1/2 (sc-514302, 1:500) from Santacruz Biotechnology, INC. Rabbit anti-ubiquitin (Z0458, 1:6000), polyclonal swine anti-rabbit IgG (P0399), polyclonal rabbit anti-mouse IgG (P0260) Horseradish Peroxidases (HRP) conjugated antibodies (1:5000) from DAKO (Glostrup, Denmark). Rabbit anti-phospho-Akt (#4060, 1:1000), and rabbit anti-Akt (#4691, 1:1000) from Cell Signaling Technologies (MA, USA). Mouse anti-phospho-Caveolin-1 (611338, 1:500) from BD (New Jersey, USA). Goat anti-SENP2 (1:6000 WB) antibodies were a gift from Dr. Ron T. Hay (College of Life Sciences, Dundee University, Dundee, UK). The specificity of the commercial antibodies has been confirmed in our previous publications (36, 37) or from laboratories that provided other antibodies.

### 2.3. Cell culture

Primary Human Skeletal Muscle Cells isolated from different skeletal muscles from adult single donors (human’s skeletal myoblasts, hSKM, A12555) from ThermoFischer Scientific. Alveolar FP-RMS: RH30; embryonal FN-RMS: RH36, RD12 a clone of RD and RD were cultured in Dulbecco’s Modified Eagle’s Medium 6046 (DMEM-high glucose, 5.5 mM, 308-340 mOs/Kg with L-Glutamine G6392) supplemented with 10% 56°C heat-inactivated Foetal Bovine/Calf Serum (FBS) and 1X 100 IU/ml of penicillin and streptomycin (Thermofischer scientific). Cells were harvested and washed with cold PBS in the presence of 0.2 M iodoacetamide. When required, different concentrations of Subasumstat (TAK-981, HY-111789, MedChemExpress) were added to the cell cultures as indicated. Stocks were prepared in DMSO, and the final working concentration did not exceed 0.01% v/v, resulting in no DMSO-induced cellular cytotoxicity.

### 2.4. Cell lysates

Cells or myotubes were lysed in RIPA buffer (25mM Tris-Cl pH 7.5, 50 mM NaCl, 0.5% NP40, 0.1% SDS, 0.5% DOC, 1mM DTT, 20mM NEM supplied with fresh Protease and Phosphatase inhibitors). A syringe with 29G needle was used to mechanically disrupt genomic DNA. Lysates were clarified at 10000 g for 15 min at 4°C. Protein concentration (DC Protein Assay Kit, Bio-Rad) was determined on the clear supernatants.

### 2.5. Immunoblotting

Desired concentrations of cell lysates were denatured for 10 min at 95°C in loading buffer (NuPage 4X, Reducing Agent 10X, Invitrogen, Carlsbad, CA) and loaded in acrylamide Bis-Tris 4-12% gradient gels (Invitrogen, Carlsbad, CA). After transfer onto PVDF membranes (Merck Millipore, Bedford, MA) for 60 min at 0.34 A, the filters were blocked in TBS-T solution (50 mM Tris-Cl, 150 mM NaCl, 0.1% Tween-20 and 5% non-fat milk, pH 7,6), and incubated with specific primary antibodies overnight at 4°C, following an incubation with the appropriate horseradish peroxidase-conjugated secondary antibodies for 1 h at room temperature. Proteins were visualized by chemioluminescence (ECL, GE Healthcare, Uppsala, Sweden), detected by the ChemiDoc MP Imaging System (Bio-Rad Laboratory, CA), and chemioluminescence intensity was analyzed only for non-saturated bands using the correspondening imaging analysis software, ImageJ version 5.0.

### 2.6. Quantitative real time PCR (qPCR)

SUMO component transcript abundance was assayed by qPCR using the primers listed in Table 1 (Sigma-Aldrich, St. Louis, MO, USA) and previously designed with PRIMER3 software (http://biotools.umassmed.edu/bioapps/primer3_www.cgi). RNAs were extracted using the GeneJET RNA purification Kit (K0731, ThermoFisher Scientific, Lithuania) and subjected to DNase I treatment (EN0521, ThermoFisher Scientific, Lithuania). The correspondent cDNAs were produced using both oligo (dT)18 and random primers using the RevertAID H Minus First strand cDNA synthesis Kit (K1632, ThermoFisher Scientific, Lithuania). qPCR reactions were performed with 100 ng of cDNA template using the SYBR Green Master Mix (A25741, Life Technologies, Carlsbad, CA, USA) in a 20 μl of final volume. The analysis was performed with QuantStudio 3 Real-Time PCR Systems, with the following cycling program: initial 50°C 2 min, denaturation 95°C 10 min, followed by 40 cycles of 95°C for 15 seconds and 60°C for 1 minute. A final step of melting curve between 65°C to 90°C, 1°C/sec temperature speed was incorporated. Relative fold change relative to the housekeeping control gene (GAPDH) was calculated as 2^-ΔCt^ (x1000) where: ΔC_t_ = C_t_(target) - C_t_(GAPDH), according to the Minimum Information for Publication of Quantitative Real-Time PCR Experiments (MIQE) guideline. We validated that similar results were obtained using two other commonly used reference genes, HPRTand Rpl38. The gene expression based on the geometric average from these genes was similar and we thus decided to keep GAPDH as reference gene. Mean and (±SD) were obtained analyses in triplicate.

### 2.7. Cell migration assay

Cell migration was quantified through wound healing assay. Cells seeded in 12-well plates (4×10^4^) formed confluent monolayers that were pre-treated for 2 hours with TAK-981 or DMSO and then wounded by scraping the cells with a 200 µl-sterile micropipette tip. After 8 and 24 hours from wound, cells were fixed with 3% paraformaldehyde (PFA)/PBS solution (20 min, 4°C) and stained with crystal violet solution (0.2% crystal violet/20% methanol/PBS) (10 min, RT). Images of wound healing were acquired at different time-points by an inverted light microscope (Olympus IX50; Olympus, Tokyo, Japan) using cellSens Software (Olympus, Tokyo, Japan). The extent of wound repair was quantified by measuring the healed area using ImageJ software. Results were presented as percentage of repaired area with respect to time 0 hours.

### 2.8. Cell proliferation assay

Cell proliferation was quantified through measurement of crystal violet incorporation. Cells were seeded in 96-well plates (2×10^3^) and treated with TAK-981 or DMSO 24 hours after seeding. At the indicated time-points cells were fixed with 3% PFA/PBS solution (20 min, 4°C) and stained with crystal violet solution (0.2% crystal violet/20% methanol/PBS) (10 min, RT). Cells were washed and resuspended in 1% sodium dodecyl sulfate (SDS)/PBS solution. Plates were shaken until complete dissolution was achieved and then absorbance was measured by reading the plate at 595 nm emission wavelength.

### 2.9. Clonogenic assay

Cells were seeded into 6-well plates in triplicate (at a density of 1000 cells/well) to evaluate clonogenic survival. After 10 days, colonies were fixed with 3% PFA/PBS solution (20 min, 4°C) and stained with crystal violet solution (0.2% crystal violet/20% methanol/PBS) (10 min, RT). Wells were washed with deionized water and pictures of colonies were taken. Then, the dye was solubilized in 1% sodium dodecyl sulfate/PBS solution. Plates were shaken until complete dissolution was achieved, and then absorbance was measured by reading the plate at 595 nm emission wavelength.

### 2.10. Cell viability assay

Cell viability was quantified by measuring the incorporation of MTT (3-[4,5-dimethyl-2-thiazolyl]-2,5-diphenyl-2H-tetrazolium bromide). Cells were seeded in 96-well plates (2×10^3^) and then treated with the indicated compounds after 24 hours. 48 hours after treatment, cells were incubated with 0.5 mg/mL MTT dissolved in cell medium (3.5 hours, 37 °C) before solubilization in DMSO. Absorbance was measured by reading the plate at 540 nm emission wavelengths.

### 2.11. Ionizing radiation treatment

Ionizing radiation treatment was performed at RT on subconfluent cells pre-treated or not with the indicated compounds for 2 hours at 4 Gray (Gy) doses using an X-ray linear accelerator (dose rate of 2 Gy/min). Post-irradiated cells were processed for clonogenic assays.

### 2.12. Bioinformatic analysis

RNA sequencing (RNA-seq) data of an RMS patient cohort of FN-RMS (n=66) and FP-RMS (n=35) samples compared to normal skeletal muscle samples (n=5) were extracted from GEO (accession number GSE108022) (2). RNA-seq data were preprocessed and normalized using the iDEP.96 tool (Prof. Xijin Ge, South Dakota State University, Brookings, SD, USA) (38), and gene expression was calculated as Log2 Counts per Million (CPM). Data were analyzed and plotted using GraphPad Prism 8.0 software (GraphPad Software, San Diego, CA, USA) and Qlucore Omics Explorer (Lund, Sweden).

### 2.13. Statistical analysis

Heteroscedastic Student’s t-tests and one-way analysis of variance (ANOVA) followed by post-hoc Tukey’s test was used to analyze the differences between two or multiple groups, respectively. *P* < 0.05 was statistically significant. Data are presented as means ± standard deviations (SDs).

## 3. Results

### 3.1 Differential SUMOylation in RMS cell lines

To assess the SUMOylation status, we analyzed global conjugation of SUMO1 and SUMO2/3 in various RMS cell lines, including alveolar FP-RMS: RH30 cells, (Figure 1, green columns); embryonal FN-RMS: RH36, RD12, RD cells, (Figure 1, red columns). These RMS cell lines were compared to normal human skeletal muscle (hSKM) cells, (Figure 1, blue columns). In hSKM, low levels of SUMO1 and SUMO2/3-conjugated proteins were observed, with a faint pattern of conjugation across a wide molecular weight range (Figure 1). In contrast, many RMS cell lines showed a marked increase in SUMO1 and SUMO2/3-conjugated proteins, particularly in RH30, RH36, RD12 and RD where a robust accumulation of high molecular weight SUMO-conjugated proteins was observed, indicative of hyper-SUMOylation (Figure 1A,C). These effects were specific to SUMO as the ubiquitin protein profile displayed a consistent pattern in all cells analyzed (Supplementary Figure 1A and B). Quantification of SUMO1 conjugates showed the highest level in RD12 cells, with nearly a threefold increase in band intensity relative to hSKM, while RH30, RH36 and RD displayed approximately two-fold increases (Figure 1B). Quantification of SUMO2/3 conjugates revealed an almost tenfold increase in RH30 compared to hSKM cells, with RH36, RD12 and RD showing significant increases of 4-to 5-fold (Figure 1D) suggesting potential heterogeneity in SUMOylation pathways within RMS subtypes. These findings indicate that SUMO conjugation is notably dysregulated in several RMS cell lines compared to normal skeletal muscle, with elevated SUMO1 and SUMO2/3 conjugates in RH30, RH36, RD12, and RD cells, suggesting that hyper-SUMOylation may contribute to the pathophysiology of RMS.

**FIGURE 1:**
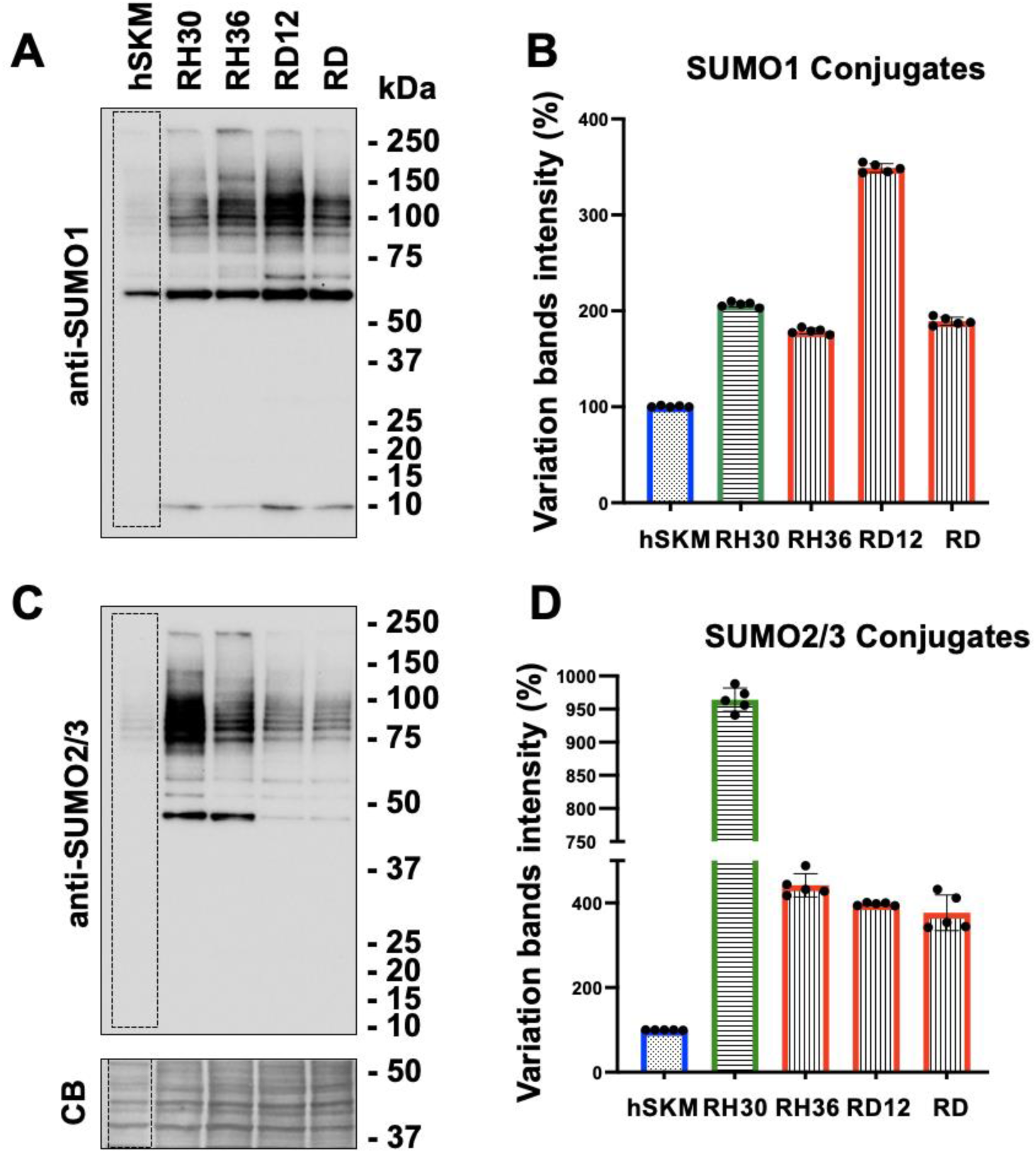
SUMOylation profile in different cell lines. Representative western blots of human skeletal muscles (hSKM), alveolar RMS (RH30) and embryonal RMS (RH36, RD12, RD) cell lysates probed with anti-SUMO1 (A) and anti-SUMO2/3 (C) antibodies showed distinct patterns of SUMO1 and SUMO2/3 conjugates. For each sample, the quantifications of the SUMO1 and SUMO2/3 conjugates were determined measuring the band intensity within the dashed areas, normalize to the correspondent total loading protein (Coomassie blue - CB) and the variations, in percentage, were referred to the hSKM as control (SUMO1 conjugates, B; SUMO2/3 conjugates, D). All data were presented as mean ± standard deviation of the average for five independent experiments.

### 3.2 Comprehensive analysis of SUMO pathway components across RMS and normal skeletal muscle cells

To investigate the differential profiles of SUMO conjugates, we performed transcriptomic analyses of SUMO moieties and SUMO-related enzymes in RMS cells from three independent cultures per cell line where each cell line exhibited a distinct SUMO transcript signature (Figure 2A,B). Principal component analysis (PCA) of SUMO transcript profiles further segregated RMS samples from control hSKM, highlighting distinct expression patterns between tumor and normal tissues.

**FIGURE 2.**
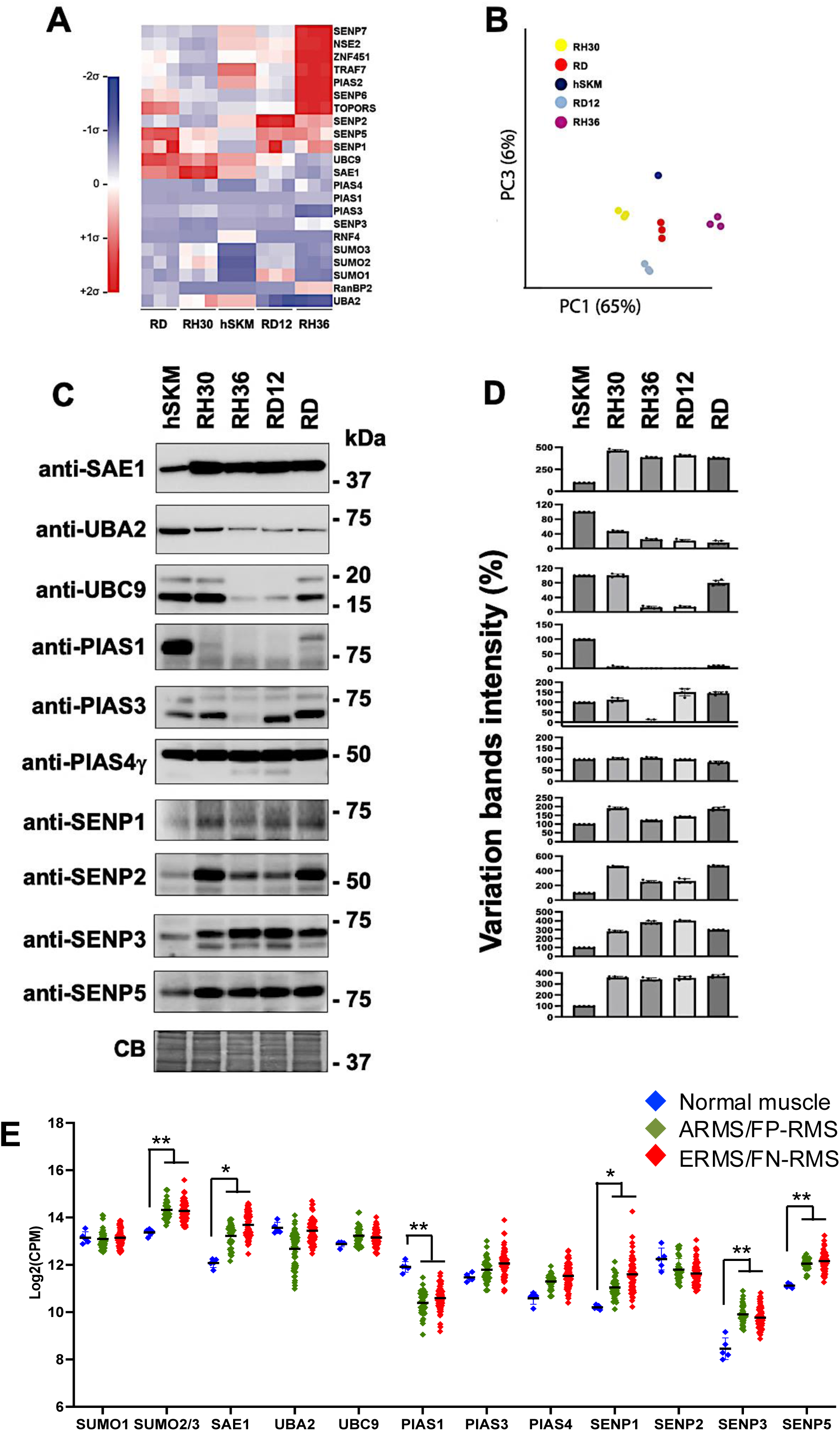
Transcriptomic analysis of the SUMO moieties (SUMO1, -2, and -3) and the SUMO enzymes from the indicated RMS cell lines. (A) Heat map shown the expression levels of all specified transcripts and normalized to GAPDH as housekeeping gene from cells cultures grown and harvested in three independent times. Values are represented as mean-centered and sigma-normalized. (B) Principal component analysis of expression data shown in A, reveals the formation of distinct clusters, indicating differences in SUMO expression signature among the RMS cells. (C) Representative western blots of the RMS cells with the indicated antibodies correspondent to validate some SUMO conjugases (SAE1, UBA2, UBC9, PIAS1, PIAS3, PIAS4γ) and deconjugases (SENP1, SENP2, SENP3, SENP5). Coomassie blue stain (CB) was used as loading control. (D) Quantification of the proteins detected on the correspondent blots. Band intensities were normalized to the correspondent total loading protein (Coomassie blue - CB) and the variations, in percentage, were referred to the hSKM as control (100%). All data were presented as mean ± standard deviation of the average for five independent experiments. (E) Expression levels of genes involved in the SUMOylation and deSUMOylation pathway across different tissue types. The scatter plot presents the log-transformed expression (Log2(CPM)) of several genes associated with the SUMOylation and deSUMOylation processes, measured in normal muscle from tissue, FN-RMS, and FP-RMS. The data points are color-coded by sample type: blue dots represent normal muscle tissue, green dots represent FP-RMS, red dots represent FN-RMS. Genes displayed along the x-axis include components of the SUMOylation pathway (e.g., SUMO1, SUMO2/3, SAE1, UBA2, UBC9) regulators such as PIAS1–PIAS4, and SUMO protease family (deSUMOylation) SENP1–SENP5. The y-axis represents the gene expression level in Log2(CPM). The plot shows variability in expression levels across the three tissue types, with notable differences in expression between normal muscle and cancerous tissues FP-RMS and FN-RMS. Statistic: **: p<0,005; *: p<0,05.

To validate these findings, cell lysates were fractionated via SDS-PAGE, and membranes were probed with antibodies specific to the SUMO pathway components E1 (SAE1, UBA2), E2 (UBC9) and E3 ligases (PIAS1, PIAS3, PIAS4γ), as well as the deconjugating enzymes SENP1, SENP2, SENP3 and SENP5 (Figure 2C,D). We observed an overall strong correlation between transcript and protein levels. Specifically, in RH30, RH36, RD12, and RD cells, SAE1 and deconjugases SENP1, SENP2, SENP3, and SENP5 were overexpressed (3-4 times higher than in hSKM), while UBA2 expression was reduced by 40-70%. UBC9 was decreased by 30-80% in embryonal RMS cells. PIAS1 was either greatly reduced or undetectable in all RMS cells, and PIAS3 was reduced only in RH36. No changes were observed in PIAS4γ expression.

To extend these findings, we performed gene expression profiling of high-throughput RNA sequencing data from 5 normal muscle from tissues and 101 RMS tumor samples (39) (Figure 2E). Correlating RMS tumor data with cultured RMS cells, we observed consistent transcriptional changes in the SUMO2/3 moieties and several SUMO enzymes. SUMO2/3 (p=0.005), SAE1 (p=0.036), SENP1 (p=0.04), SENP3 (p=0.001), and SENP5 (p=0.002) were upregulated, while PIAS1 (p=0.004) was downregulated, while UBA2 (p=0.51), showed a trend of downregulation in both tumor samples and RMS cell lines. These results suggest that alveolar and embryonal RMS cells serve as suitable models for further investigation.

### 3.3 Heat shock stress response to RMS and control cells

The SUMOylation pathway is a critical cellular sensor that detects physiological changes and responds by modulating SUMO-conjugated proteins (40). In the above experiments, we identified variations in SUMO enzyme levels among RMS cells. To explore how SUMOylation responds to stress, we here subjected cells to 43°C heat shocks for 30 minutes, followed by recovery at 37°C for 2 hours. In human control muscle, a typical SUMOylation response to heat shock was observed. Western blot analyses again reveal that RMS cells exhibit significantly higher levels of SUMO1- and SUMO2/3-conjugated proteins compared to normal skeletal muscle. During the heat shock, SUMOylated proteins accumulated and after the recovery period SUMOylation returned to baseline levels (Figure 3A,B). Similar patterns were found for RH36 and RD12 cells. In contrast, RH30 and RD cells maintained elevated levels of SUMOylated proteins even after the recovery period (Figure 3B,D). These results provide functional support for the impaired SUMOylation process in these cells due to the alteration of the SUMO enzymes and enhanced SUMOylation response in cancer cells, suggesting its potential role in stress responses and possibly in cancer progression or survival under adverse conditions.

**FIGURE 3.**
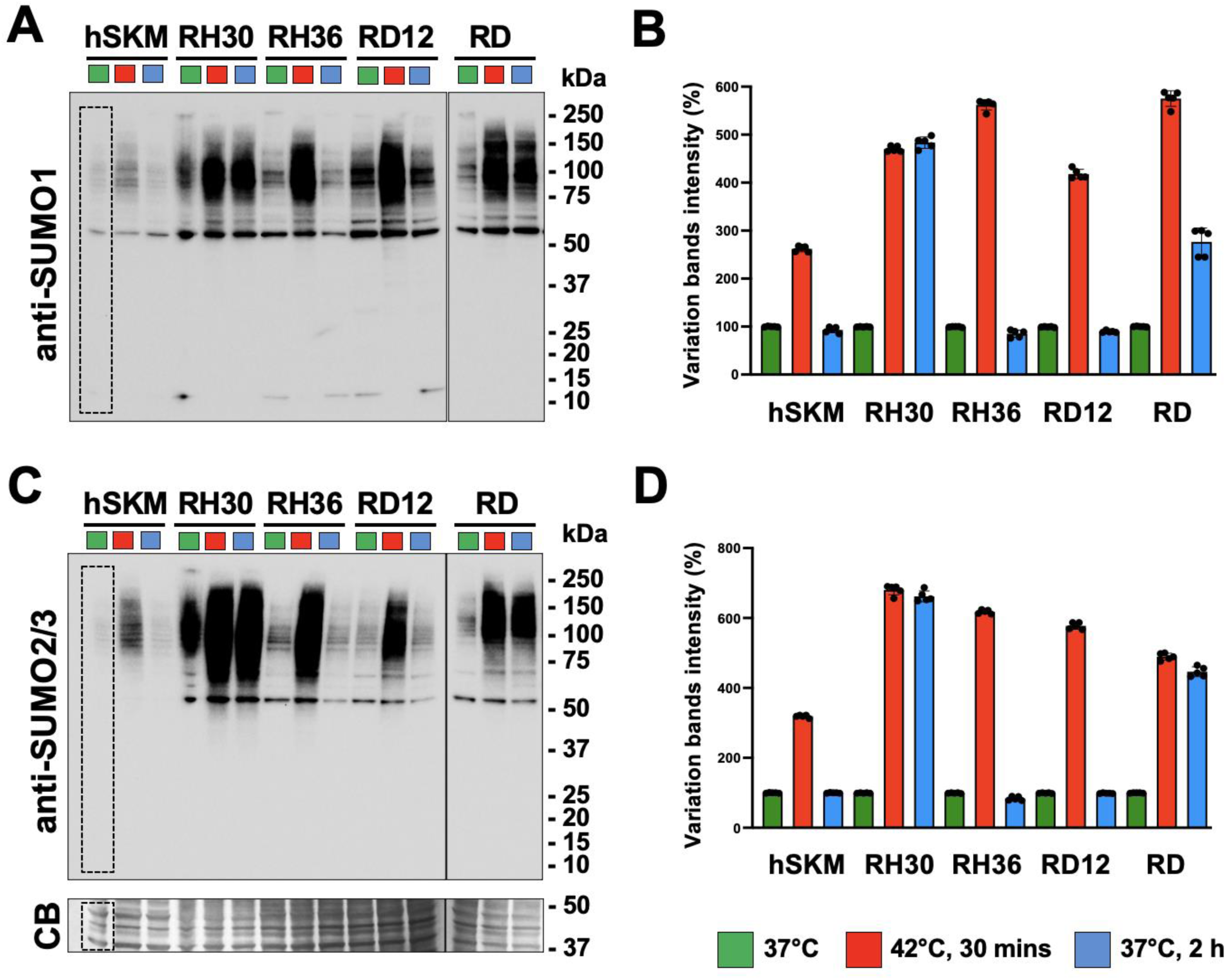
Heat shock and recovery treatments of RMS cells. Lysates of hSKM and RMS cells grown at 37°C (green color), at 42°C for 30 minutes (red color) and then replaced at 37°C (blue color) for 2 hours were separated in SDS-Page gels and membranes probed with anti-SUMO1 (A) and anti-SUMO2/3 antibodies (C). Band intensity quantification of the SUMO1 (B) and SUMO2/3 conjugates were determined within the dashed areas, normalize to the total loading protein (Coomassie blue - CB) and the variations, in percentage, were referred to the correspondent sample grown at 37°C (green color). All data were presented as mean ± standard deviation of the average for five independent experiments.

### 3.4 TAK-981 inhibits SUMO conjugation and reduces cell migration, proliferation and colony formation of RMS cells

TAK-981 is a potent and highly selective first-in-class SUMOylation inhibitor with demonstrated anticancer efficacy that it is currently undergoing clinical trials for solid tumors (35) (41) (42). RMS is a soft tissue cancer, and both alveolar and embryonal RMS cells display an accumulation of SUMOylated proteins compared to human myoblasts. We first evaluated the effects of TAK-981 on control hSKM cells and different RMS cell lines. After 4 hours of treatment, a dose-dependent reduction in SUMO1 and SUMO2/3 conjugates was observed, in RH36, RD12, RD, and hSKM cells starting at 10 nM, with reductions between 80 − 60% of total SUMO conjugates, whereas RH30 cells remained unaffected, even at 100 nM (Supplementary Figure 2). When extending exposures to 72 hours, complete SUMO deconjugation was achieved in RH36 and RD12 cells with 1 nM TAK-981, while in RD and hSKM cells, SUMOylation was completely blocked at 10 nM. RH30 cells became responsive, showing a 25% reduction at 1 nM TAK-981 and a 50% inhibition at 100 nM; however, RH30 cells remained least the sensitive among the tested cell lines (Figure 4A,B).

**FIGURE 4.**
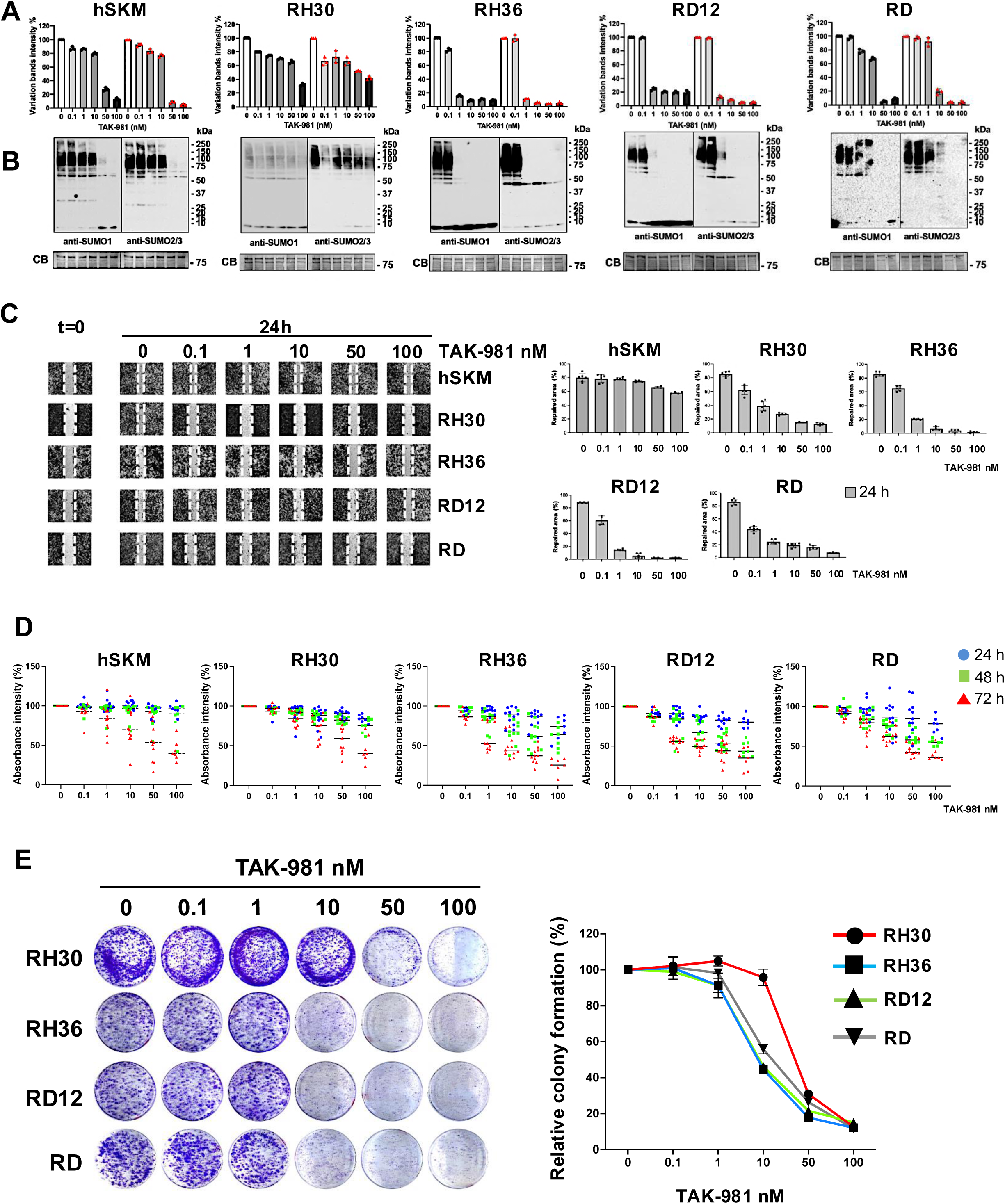
Impact of TAK-981 on SUMOylation, cell migration, proliferation and colony formation in RMS and control cells. Quantification of global SUMOylation in hSKM (human skeletal muscle) cells and four RMS cell lines (RH30, RH36, RD12, and RD) following treatment with increasing concentrations of TAK-981 (0, 0.1, 1, 10, 50, and 100 nM). Data are expressed as the percentage of variation in band intensity compared to control (0 nM). Black symbols referred to SUMO1 and red symbols referred to SUMO2/3 conjugates (A). Western blot of SUMO1 and SUMO2/3-conjugated proteins in hSKM, RH30, RH36, RD12, and RD cells treated with TAK-981. The blots demonstrate the drug effects on SUMO1 and SUMO2/3 conjugation. CB refers to Coomassie Blue staining, used as a loading control. Average and standard deviation are calculated from three independent experiments (B). Representative images of the wound healing assay at time 0 and 24 hours for hSKM and RMS cell lines (RH30, RH36, RD12, and RD) treated with TAK-981 at the indicated concentrations. Images show the progression of wound closure, with the dashed lines marking the wound edges at t=0. Cell migration percentage was quantified 24 hours after treatment. Average and standard deviations are calculated for n=6 independent experiments (C). Cell proliferation was evaluated by measuring absorbance intensity in hSKM, RH30, RH36, RD12, and RD cells treated with increasing concentrations of TAK-981 (0, 0.1, 1, 10, 50, and 100 nM) for 24 (blue symbols), 48 (green symbols), and 72 (red symbols) hours. The colored dots represent the individual data points for the different time points, and bars indicate the average of n=12 independent experiments (D). Colony Formation Assay: RMS cell lines (RH30, RH36, RD12, and RD) were treated with increasing concentrations of TAK-981 (0, 0.1, 1, 10, 50, and 100 nM). Representative images of colonies stained with crystal violet are shown for each cell line and treatment condition. The graph on the right shows quantification of colony formation relative to control (0 nM TAK-981), with data expressed as a percentage. Average and standard deviation are calculated from three independent experiments (E).

Next, we evaluated the functional consequences of SUMO inhibition. In wound healing assays, all cells exhibited delayed repair compared to DMSO-treated controls with a notable effect observed as early as 8 hours after treatment (Supplementary Figure 3A). After 24 hours, control hSKM cells almost fully repaired the wound, with only 100 nM TAK-981 impairing repair by ∼20%. In contrast, RMS cells exhibited significantly impaired migration. RD cells showed >50% reduction in migration already at 0.1 nM, while sensitivity was slightly lower for RH36, RD12, and RH30 cells (Figure 4C).

After, we investigated the effect of TAK-981 on cell proliferation after 24, 48, and 72 hours of treatment using crystal violet assays (Figure 4D). A reduction of nearly 30% in cell proliferation was evident after 24 hours and 50% after 72 hours in RH36, RD12, and RD cells treated with 10 nM TAK-981. In contrast RH30 and hSKM cells were less affected and a comparable reduction in proliferation was only observed after 72 hours with 50 nM TAK-981. Control hSKM cells exhibited minimal sensitivity to TAK-981, suggesting selective inhibition in cancerous cells.

To explore the antitumorigenic effects of TAK-981 in both alveolar and embryonal cells, we conducted colony formation assays. A significant reduction of 50-60% in colony formation was observed in embryonal cell lines (RH36, RD12, and RD) at 10 nM TAK-981, whereas alveolar cells (RH30) were less sensitive and required five-fold higher concentrations to show a significant response (Figure 4E). Functionally, TAK-981 impairs cell migration, proliferation, and colony formation in RMS cells, with embryonal RMS cells showing greater sensitivity than alveolar RMS cells. Minimal effects on control hSKM cells suggest TAK-981’s selective anticancer potential.

### 3.5 TAK-981 sensitizes RMS cells to chemotherapy but not to radiotherapy

We next examined whether TAK-981 could exert a synergistic effect when combined with other chemotherapeutic agents commonly used in RMS therapy, specifically actinomycin D and doxorubicin. Importantly, effects of 10 nM TAK-981 were additive with 0.5 nM actinomycin D or 0.5 μM doxorubicin for RH30 (p=0,0001; p=0,002), RH36 (p=0,0001; p=0,0001), and RD12 (p=0,002; p=0,002), (Figure 5A). For RD, data even suggested synergy between SUMO inhibition and actinomycin D (p<0,0001), and doxorubicin (p=0,002).

**FIGURE 5.**
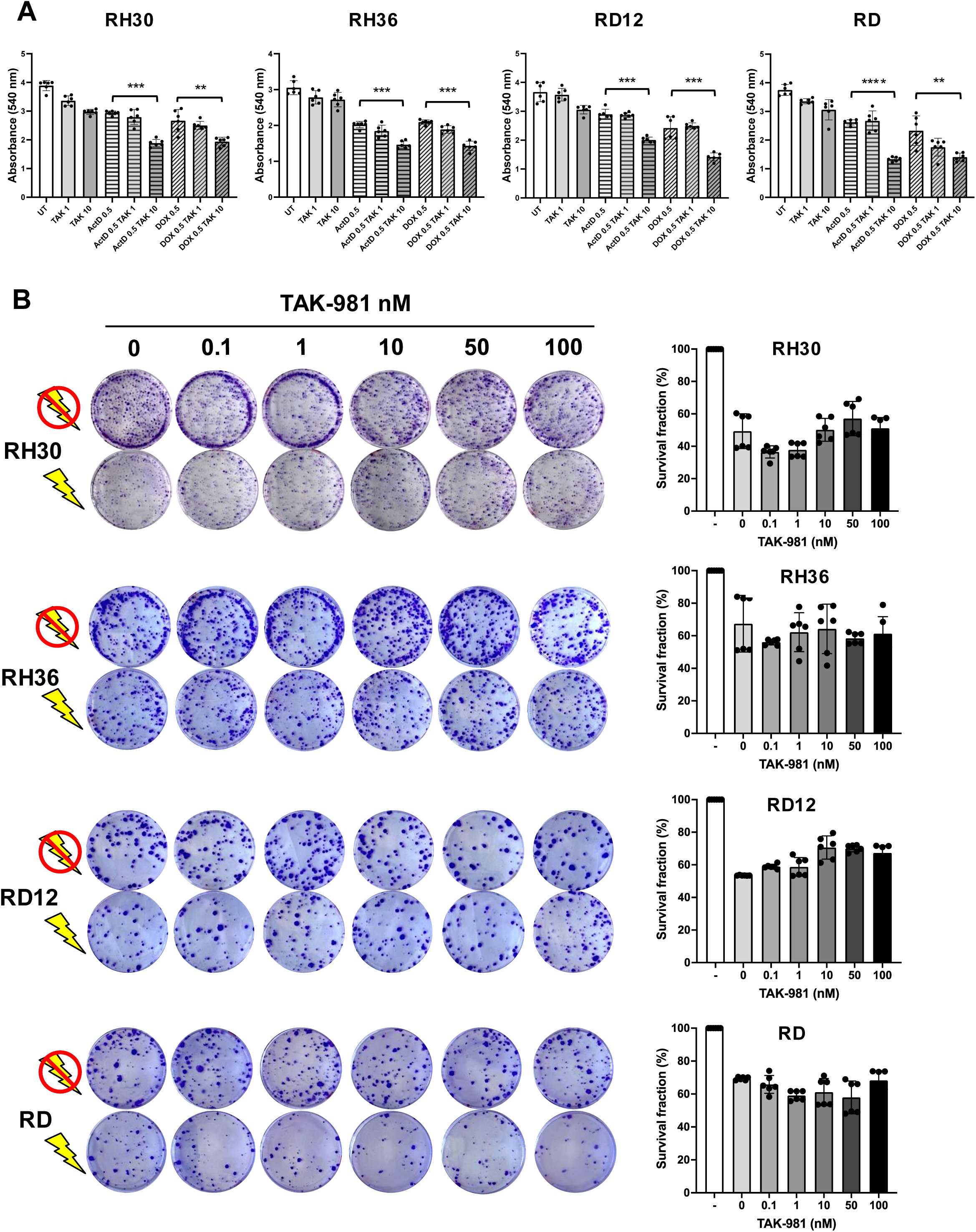
TAK-981 enhances the efficacy of chemotherapy when combined with other drugs but does not influence radiosensitivity. RH30, RH36, RD12, and RD were treated with TAK-981 alone or in combination with chemotherapeutic agents (Actinomycin D, ActD, or Doxorubicin, DOX). Bar graphs show cell viability measured by absorbance at 540 nm under different treatment conditions: untreated (UT), TAK-981 (1 nM, 10 nM), ActD (0.5 nM), DOX (0.5 µM), and combination treatments. Each bar represents mean ± SD from n=6 replicates, and results indicate that TAK-981 alone and in combination with ActD or DOX reduces cell viability in all tested RMS cell lines (B). Clonogenic survival assay of RMS cell lines after TAK-981 treatment with and without irradiation. Colony formation in RMS cell lines (RH30, RH36, RD12, and RD) treated with increasing concentrations of TAK-981 (0, 0.1, 1, 10, 50, 100 nM) both in the presence (lightning symbol) and absence (no symbol) of radiation. Cells were stained with crystal violet to visualize colonies. Quantification of the survival fraction (percentage of surviving colonies) after treatment with TAK-981 and irradiation in RH30, RH36, RD12, and RD cell lines. Bars represent the mean survival fraction ± SD across different TAK-981 concentrations of n=6 independent experiments. Statistic: ****: p<0,0001; ***: p=0,0001; **: p=0,002.

To investigate whether TAK-981 also influences radiosensitivity, we treated the RMS cell lines with varying concentrations of TAK-981, followed by irradiation (Figure 5B). Colony formation assays revealed no significant differences in survival fractions between cells treated with TAK-981 alone and those treated with TAK-981 followed by irradiation at any concentration tested. These results indicate that TAK-981 does not affect radiosensitivity in RMS cells. These data suggest that TAK-981 is a potent inhibitor of SUMO conjugation that demonstrates antitumor activity both as a monotherapy and in combination with other chemotherapeutic agents in both alveolar and embryonal RMS subtypes.

### 3.6 TAK-981 inhibits AKT, ERK, and CAV1 phosphorylation

Since AKT, ERK, and caveolin-1 (CAV1) are key regulators of cell migration and proliferation (43) (44), we investigated the impact of TAK-981 on their activation states (45) (46) (47). TAK-981 caused a significant reduction in phosphorylated AKT (p-AKT), with a >50% decrease in RH36, RD12 and RD cells at 1 nM (Figure 6A). RH30 and hSKM cells were less sensitive with relevant reduction of AKT phosphorylation being only apparent at 50 nM. Total AKT levels were unaffected. Similarly, total ERK levels remained stable, while p-ERK was reduced by >80% in RD12 and hSKM cells at 10 nM, and in RH36 and RD cells at 50 nM (Figure 6B). In RH30 cells, a 70% reduction in p-ERK required 100 nM of TAK-981. Interestingly, ERK phosphorylation was most sensitive in hSKM and RD12 cells, indicating that the sensitivity of SUMO effects on the different downstream pathways is cell type specific.

**FIGURE 6.**
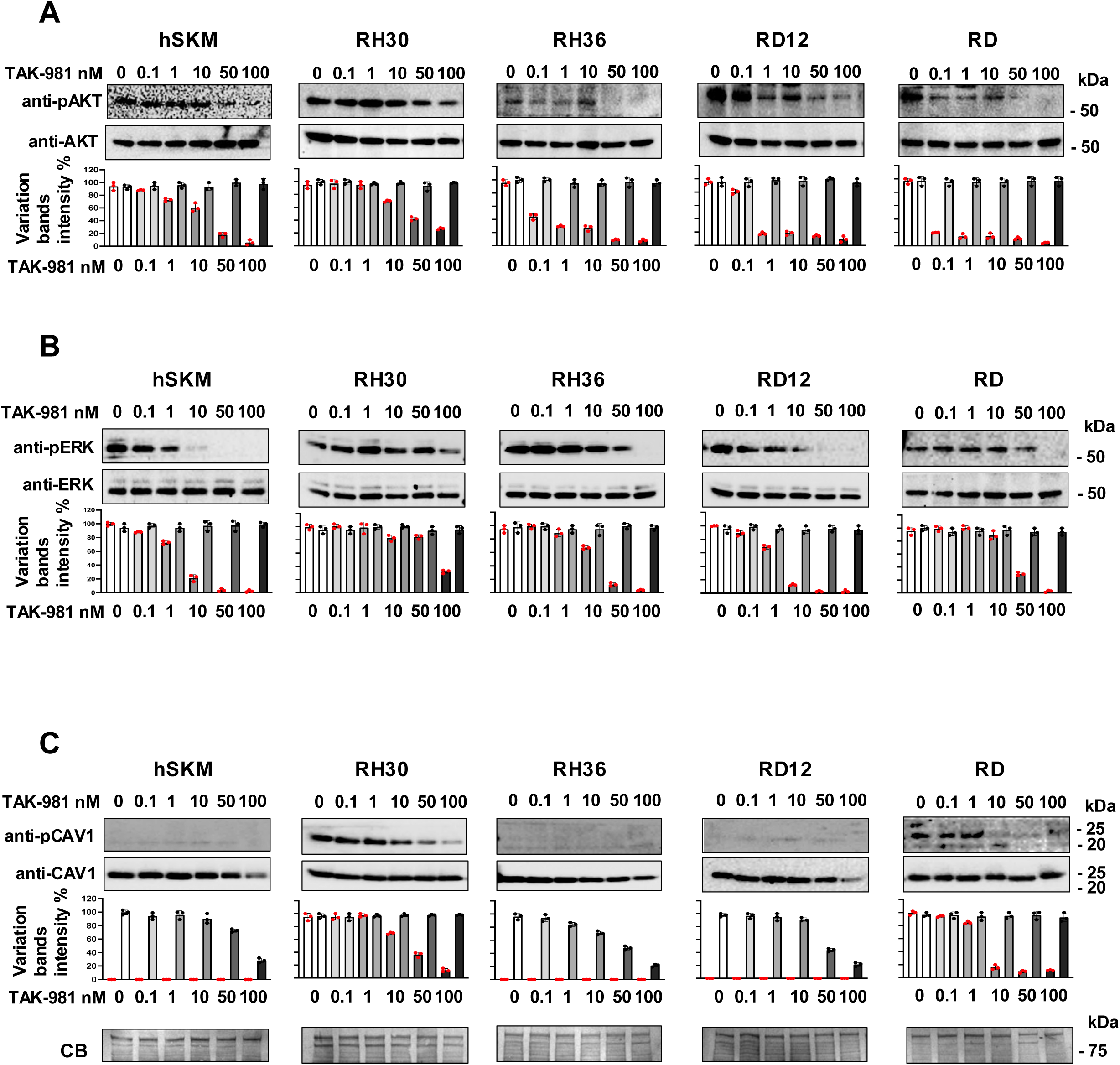
TAK-981 inhibits key signaling pathways in RMS cell lines after 72h of treatment. Western blot analysis showing the dose-dependent inhibition of pAKT (phosphorylated AKT) levels upon treatment with TAK-981 in a panel of RMS cell lines (RH30, RH36, RD12, RD) and control hSKM (human skeletal muscle cells). Cells were treated with TAK-981 at concentrations of 0, 0.1, 1, 10, 50, and 100 nM. Blots were probed with antibodies against pAKT (Ser473) and total AKT. The intensity of pAKT bands is progressively reduced in most cell lines, indicating the TAK-981 ability to downregulate AKT activation. The corresponding histograms below the blots depict the quantification of pAKT (red symbols) and the total AKT (black symbols), showing the relative levels at different concentrations of TAK-981 and referred to the vehicle (0 nM) treatment (A). Western blots showing the effect of TAK-981 on ERK phosphorylation (pERK) in the same panel of cell lines. Antibodies against pERK and total ERK were used. TAK-981 treatment leads to a dose-dependent reduction in pERK levels across most cell lines, as seen in the blot intensities. The quantification (bar graphs below) illustrates the pERK levels (red symbols) and the total ERK (black symbols), showing the relative levels at different concentrations of TAK-981 and referred to the vehicle (0 nM) treatment (B). Western blots demonstrating the effect of TAK-981 on phosphorylated caveolin-1 (pCAV1) and total CAV1 in RMS cell lines and hSKM cells. The cells were treated with the same doses of TAK-981 (0, 0.1, 1, 10, 50, 100 nM). Quantification of the phosphorylated CAV1 (pCAV1) (red symbols) and total CAV1 (black symbols) is shown in the bar graphs below (C). The loading control for the Western blots is shown as Coomassie Blue staining of total protein. The blots are a representation of n=3 experiments performed independently.

Phosphorylated CAV1 (p-CAV1) was only detected in RH30 and weakly in RD cells. In RD, p-CAV1 was undetectable at 10 nM TAK-981, while in RH30, >60% reduction was observed with 50 nM. In RH36 and RD12 cells, where p-CAV1 was absent, total CAV1 protein levels decreased by ∼50% at 50 nM TAK-981, and by the same percentage in hSKM cells at 100 nM. Thus, TAK-981 effectively inhibited the phosphorylation of AKT, ERK, and CAV1 in a dose-dependent manner in different RMS cell lines with subtype-specific sensitivity profiles, demonstrating its potential as a therapeutic agent targeting multiple signaling pathways.

## 4. Discussion

This study underscores the pivotal role of SUMOylation in defining the pathophysiology of RMS and highlights its broader implications for tumor behavior, stress responses, and therapeutic intervention. It is reasonable to infer that elevated levels of SUMO, and E1, E2 or E3 enzymes lead to increased SUMOylated proteins. Conversely, higher levels of deSUMOylating enzymes may enhance the reduction of SUMO conjugates. Since maintaining the SUMOylation equilibrium is crucial for all cells, alteration in the SUMO pathway components has been described as a mechanism driving neoplastic diseases (21).

Our findings reveal significantly elevated levels of SUMO1 and SUMO2/3 conjugates in RMS cell lines, particularly in RH30 (Alveolar/FP-RMS) and RH36, RD12, and RD (Embrional/FN-RMS) cells, compared to normal skeletal muscle (hSKM). This hyper-SUMOylation likely drives tumor progression through its effects on key cellular processes such as DNA repair, transcription, and cell cycle regulation (20) (40).

Transcriptomic analyses reveal that the varying levels of SUMOylation among RMS cells subtypes are attributable to differential expression of SUMO enzymes. This heterogeneity underscores subtype-specific dependencies on the SUMOylation machinery. Alveolar and embryonal RMS cells appear to leverage hyper-SUMOylation to adapt to distinct oncogenic pressures, emphasizing the role of SUMOylation in shaping tumor phenotype and behavior.

The overexpression of SAE1, a component of the SUMO E1 enzyme and a potential biomarker of this group of cancer, along with SUMO maturation-specific proteases SENP1 and SENP3(17), was consistent across all RMS cells and tumor samples analyzed. These findings implicate these enzymes as primary drivers of hyper-SUMOylation.

Together with its role in tumor progression, altered SUMOylation impacts cellular stress responses. For example, RH30 and RD cells exhibit an aberrant SUMOylation response to heat shock, maintaining elevated levels of SUMO-conjugated proteins even after heat shock recovery, in stark contrast to normal skeletal muscle. This persistent SUMOylation likely enhances RMS cells’ ability to survive under stress, facilitating tumor adaptation and progression. The differential stress responses among RMS subtypes highlight the complexity of SUMO-dependent regulatory mechanisms and their role in tumor resilience.

The centrality of SUMOylation in RMS pathophysiology makes the SUMO pathway a compelling therapeutic target. TAK-981, a potent SUMOylation inhibitor that binds to SAE1, effectively impaired tumorigenic processes such as proliferation, migration, and colony formation in RMS cells, demonstrating its potential as a monotherapy or adjunct in RMS treatment. Interestingly, the differential sensitivity of RMS subtypes to TAK-981 highlights the complexity of SUMOylation’s role in tumor biology. Embryonal RMS cells showed greater sensitivity to TAK-981 than alveolar RMS cells, with RH30 cells exhibiting partial resistance. These findings necessitate further exploration of subtype-specific SUMO dependencies to optimize therapeutic strategies. A key finding of this study is the ability of TAK-981 to enhance the efficacy of commonly used chemotherapeutic agents, such as actinomycin D and doxorubicin, in RMS cell lines, while its lack of impact on radiosensitivity highlights a nuanced limitation.

This result aligns with previous reports showing that TAK-981 amplifies the effect of rituximab in promoting innate immune responses (48) and synergizes in combination with 5-azacytidine in the preclinical acute myeloid leukemia treatment(49). The observed synergistic effect, where TAK-981 in combination with these drugs resulted in greater inhibition of cell proliferation compared to monotherapy, underscores its potential as an adjunct to chemotherapy in RMS treatment. This is particularly relevant for RMS, where treatment resistance and recurrence remain significant clinical challenges. The synergy observed with chemotherapy may arise from SUMO’s influence on transcriptional and DNA repair pathways, which are differently involved in radiation-induced damage responses. Indeed, cells undergoing active proliferation are typically the most susceptible to radiation-induced damage. Consequently, a drug that inhibits cell proliferation may not necessarily enhance sensitivity to radiotherapy.

The impact of SUMOylation inhibition extends beyond direct tumorigenic effects to the modulation of key signaling pathways. TAK-981’s ability to reduce phosphorylation of AKT, ERK, and caveolin-1 underscores its broad regulatory role in cellular survival and proliferation pathways. This multi-faceted mechanism suggests that targeting SUMOylation not only disrupts tumor growth but also alters the signaling networks that sustain oncogenic phenotypes.

However, the overarching dysregulation of SUMO enzymes (e.g., overexpression of SAE1, SENPs, and reduced UBA2) presents additional avenues for genetic intervention. These molecular imbalances correlate with poor survival outcomes, reinforcing the prognostic and therapeutic value of targeting SUMO components.

In this study, we provided solid evidence that SUMOylation emerges as a cornerstone of RMS pathophysiology, orchestrating tumor progression, stress response, and therapeutic resistance. The dysregulation of SUMOylation in RMS, combined with its subtype-specific nuances, highlights its importance as both a biological hallmark and a therapeutic target.

While TAK-981 demonstrates promise as a SUMOylation inhibitor with potent anticancer activity (41) (42) (50), it is recently being tested in combination with other drugs to treat people who have select advanced or metastatic solid tumors of non-squamous non-small-cell lung cancer (NSCLC) or microsatellite-stable colorectal cancer (MSS-CRC), (Clinical trial information: NCT04381650). However, within the range of TAK-981 concentrations (0.1–100 nM) employed in our in vitro experiments, the maximum concentration of TAK-981 achieved at the recommended phase 2 dose in humans exceeds 1 µM (51).

For RMS, further investigation into the molecular intricacies of the SUMO pathway and mechanisms of drug resistance is crucial. Our findings support the integration of SUMO-targeted therapies with existing treatment modalities, emphasizing the need for subtype-specific approaches. By advancing our understanding of SUMOylation in RMS, we further aim to characterize and identify the SUMO1 and SUMO2/3 conjugates, and develop more effective and personalized interventions, addressing the persistent challenges of treatment resistance and recurrence in this aggressive malignancy.

### Conclusions

This study highlights SUMOylation as a critical driver of RMS pathophysiology, influencing tumor progression, stress response, and therapeutic resistance. Hyper-SUMOylation and dysregulated SUMO enzymes underscore subtype-specific vulnerabilities. TAK-981 shows promise in impairing tumorigenic processes and enhancing chemotherapy, highlighting SUMOylation as a therapeutic target. Further research into SUMO pathways and resistance mechanisms is vital for optimizing RMS treatments.

## Supporting information

Uncropped blots

## Declaration sections

### Consent for publication

Not applicable.

### Ethics, Consent to Participate, and Consent to Publish declarations

Not applicable.

### Availability of data and Materials

The corresponding author will provide original data and materials upon request.

### Author’s contributions

S.C., E.V.C., S.G. performed the experiments. V.M.L. performed bioinformatic analysis and calculations. S.C., S.G. analyzed the data. All authors contributed to the final version of the manuscript. The study was conceived and supervised by SG.

## Abbreviations

PTM(s): Post-translational modification(s)
SENP(s): Sentrin-specific proteases
SUMO: Small Ubiquitin-like Modifier
SUMO E1: SUMO-activating enzyme E1
SAE1/2: SUMO activating enzyme subunits 1/2
SUMO E2: SUMO-conjugating enzyme E2
SUMO E3: SUMO ligase E3
UBC9: Ubiquitin conjugating enzyme 9.
PIASs: Protein inhibitor of activated STAT superfamily.
pAKT: Phosphorylated AKT.
pERK: Phosphorylated ERK.
pCAV1: Phosphorylated Caveolin-1.
hSKM: human Skeletal Muscle Cells.
PML: Promyelocytic Leukemia Protein.

## Supplementary figures

**Supplementary Figure 1.**
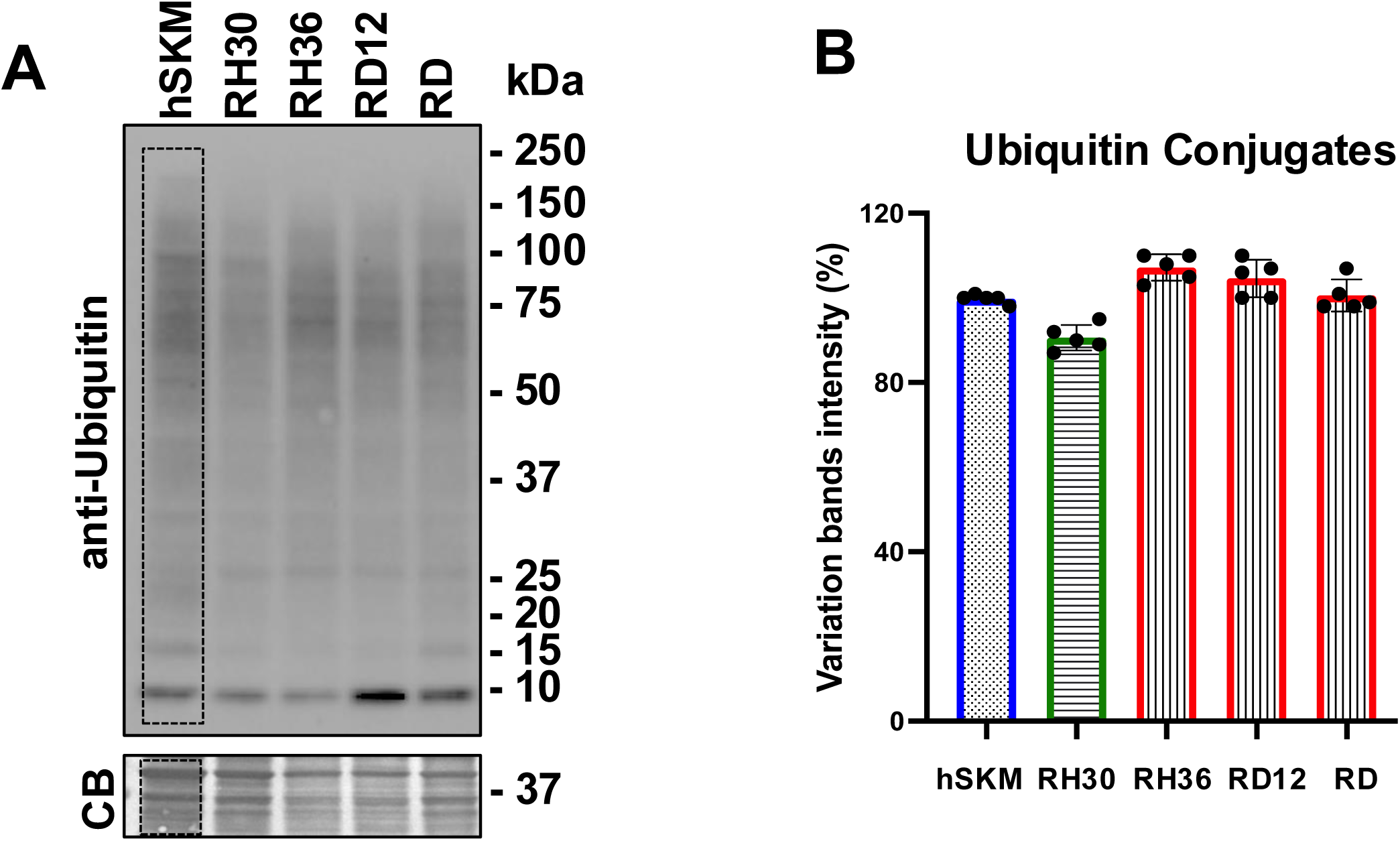
Analysis of ubiquitin conjugates expression in RMS cell lines and control. (A) Immunoblot analysis of ubiquitin conjugates in RMS cell lines (RH30, RH36, RD12, RD) and human skeletal muscle (hSKM) used as control. Proteins were separated by SDS-PAGE and probed with anti-ubiquitin antibody. A Coomassie Blue (CB) stain of total protein is shown as a loading control. Molecular weight markers (kDa) are indicated on the right. (B) Quantification of ubiquitin conjugates intensity relative to the hSKM control. Data are presented as mean ± SEM from at least three independent experiments. The variation in band intensity is expressed as a percentage relative to hSKM.

**Supplementary Figure 2.**
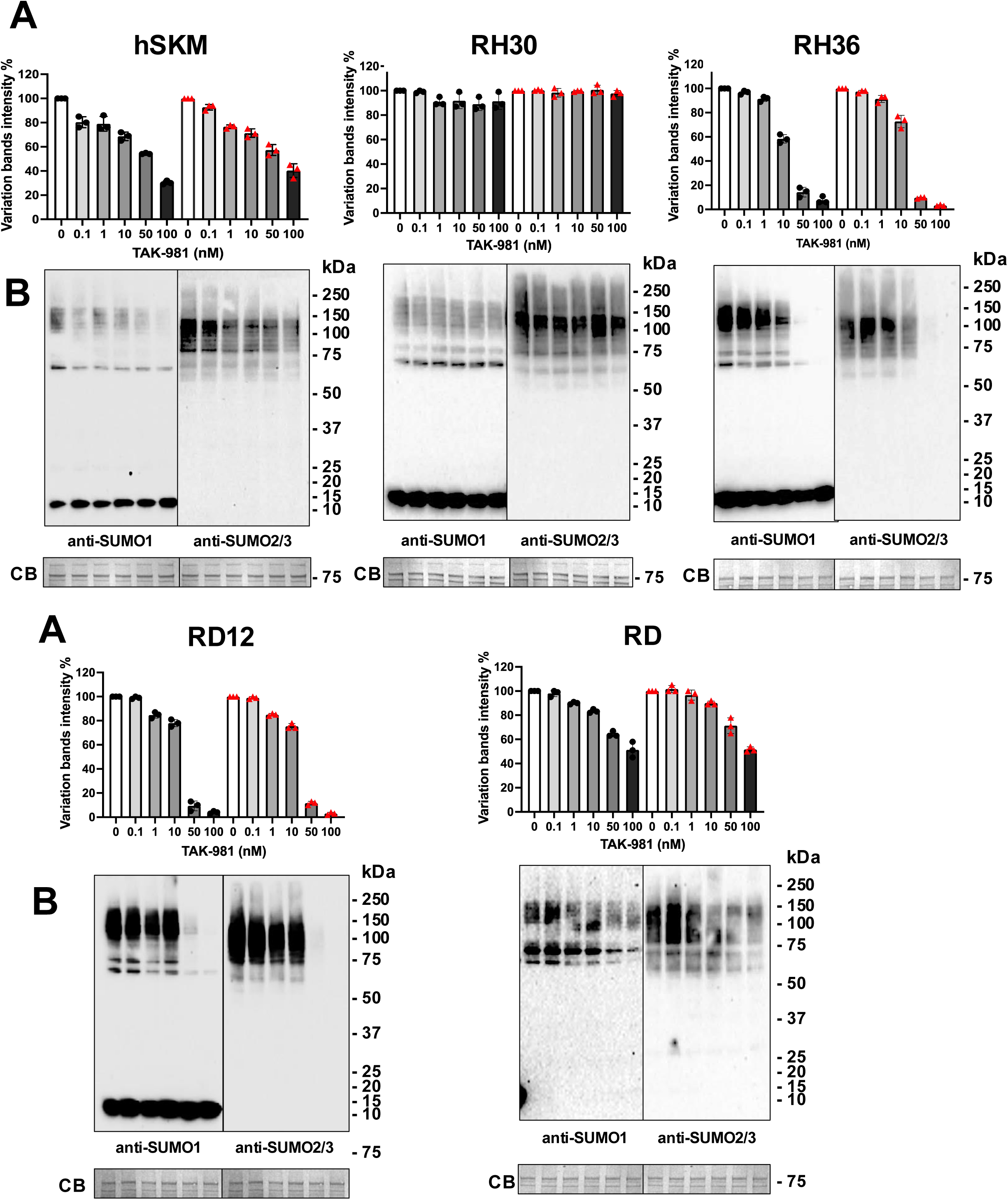
Dose-dependent inhibition of SUMOylation by TAK-981 in RMS cell lines and human skeletal muscle (hSKM) control after 4 hours. Western blot analysis of SUMO1 and SUMO2 conjugates in hSKM and RMS cell lines (RH30, RH36, RD12, RD) treated with varying concentrations of the SUMOylation inhibitor TAK-981 (0, 0.1, 1, 10, 50, 100 nM) for 4 hours. Blots were probed with anti-SUMO1 and anti-SUMO2 antibodies to detect SUMO-conjugated proteins. Coomassie Blue (CB) staining is provided as a loading control. Molecular weight markers (kDa) are indicated on the right. Top panels: Quantification of SUMO conjugate intensity in hSKM and RMS cell lines following TAK-981 treatment. Band intensity is expressed as a percentage relative to untreated control (0 nM). Data are shown as mean ± SEM from three independent experiments. Bottom panels: Representative immunoblots showing the dose-dependent reduction of SUMO1 and SUMO2 conjugates across the cell lines. Molecular weight markers (kDa) are indicated.

**Supplementary Figure 3.**
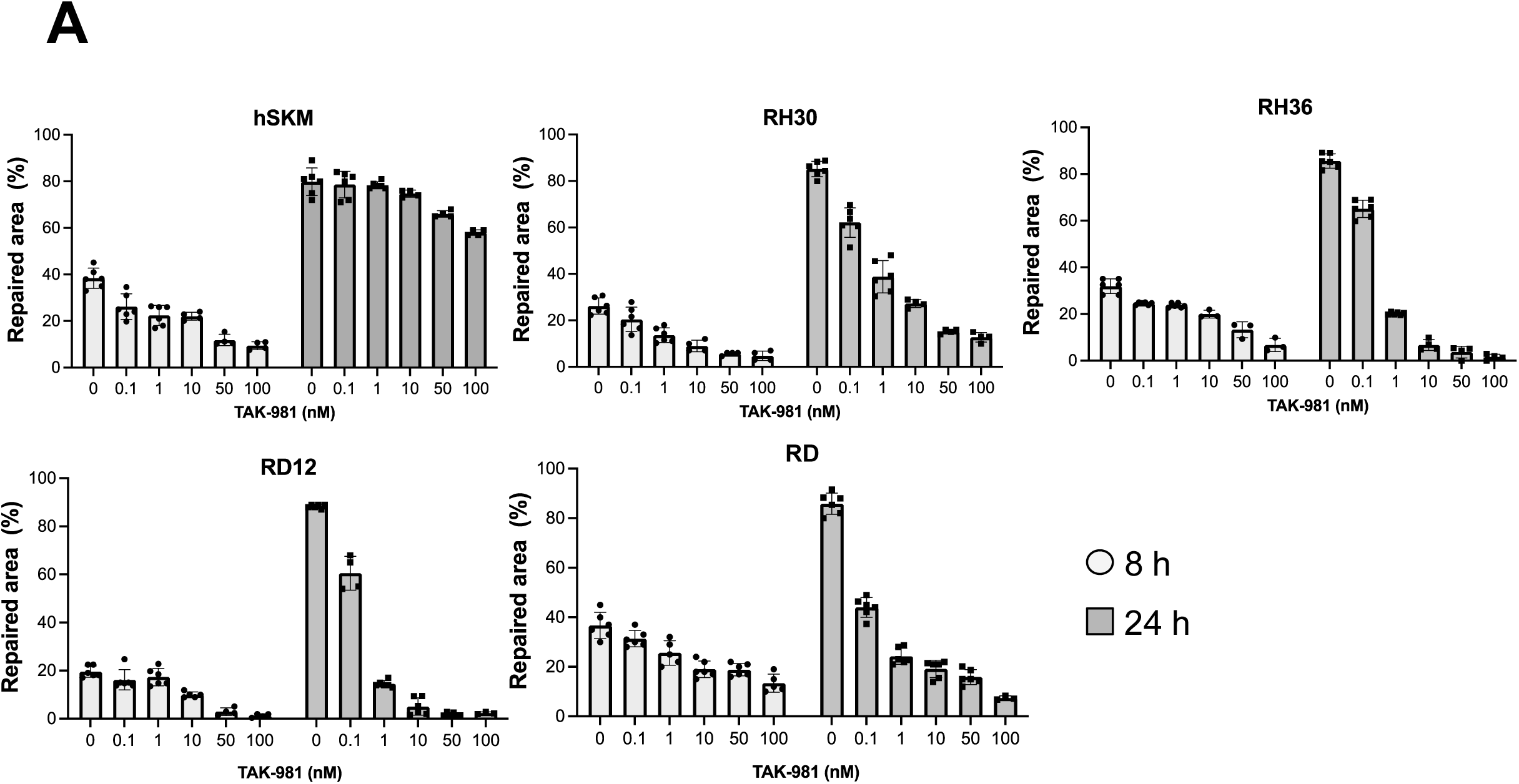
Effect of TAK-981 on cell migration in RMS cell lines and human skeletal muscle (hSKM) control. (A) Quantification of wound healing (WH) assay measuring the cell migration ability of hSKM and RMS cell lines (RH30, RH36, RD12, RD) following treatment with increasing concentrations of TAK-981 (0, 0.1, 1, 10, 50, 100 nM) for 8 and 24 hours. The percentage of the repaired area was calculated relative to the initial wound area at 0 hours. Data represent mean ± SEM from three independent experiments.

