## Supplementary material for "Synthetic inhibition of the SUMO pathway by targeting the SAE1 component via TAK-981 compound impairs tumor growth and chemosensitizers embryonal and alveolar rhabdomyosarcoma cell lines": Uncropped blots

### Slide 1
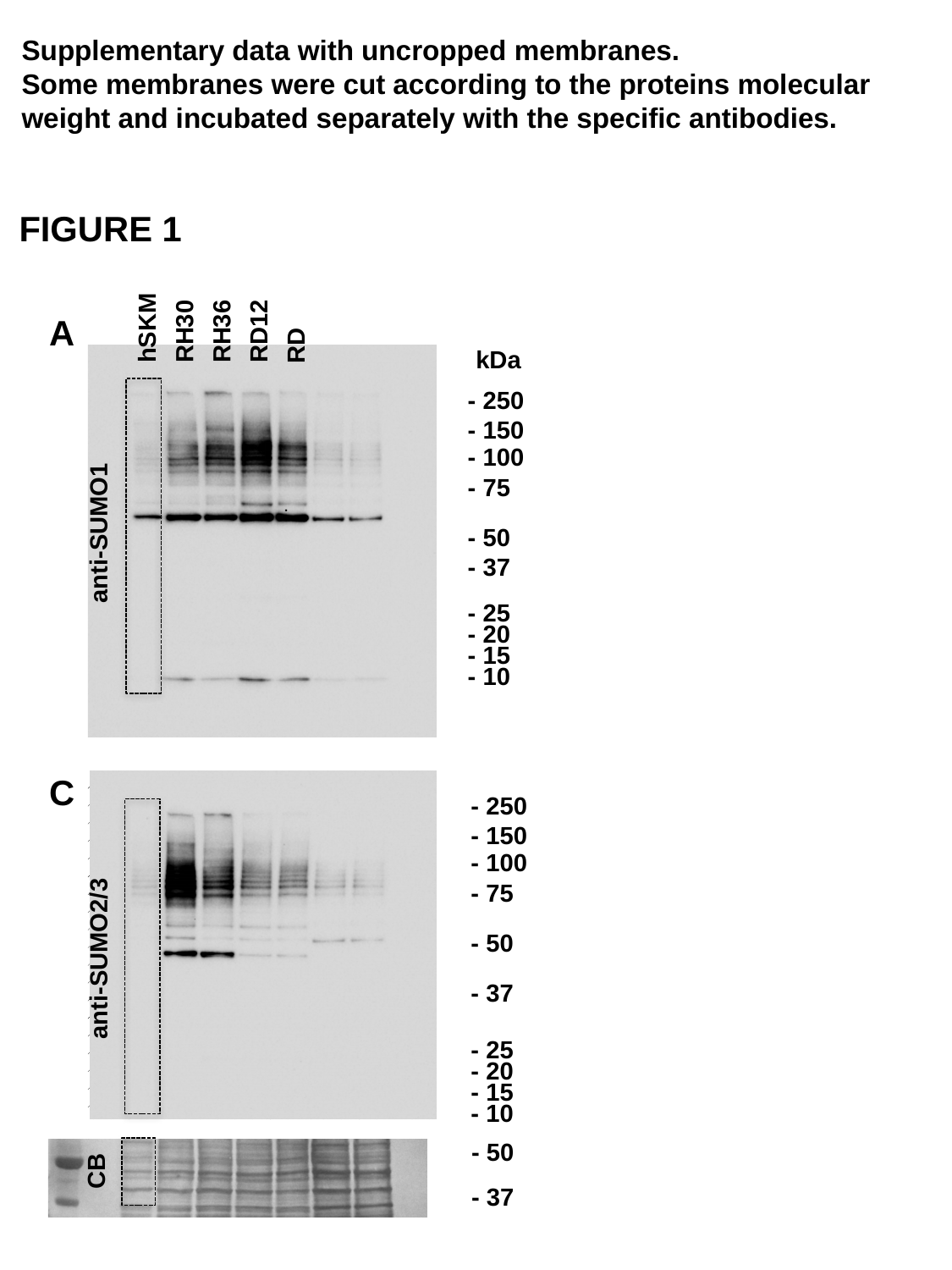

Supplementary data with uncropped membranes.
Some membranes were cut according to the proteins molecular weight and incubated separately with the specific antibodies.
FIGURE 1
RD
hSKM
RH30
RH36
RD12
A
kDa
- 250
- 150
- 100
- 75
anti-SUMO1
- 50
- 37
- 25
- 20
- 15
- 10
C
- 250
- 150
- 100
- 75
- 50
anti-SUMO2/3
- 37
- 25
- 20
- 15
- 10
- 50
CB
- 37

### Slide 2
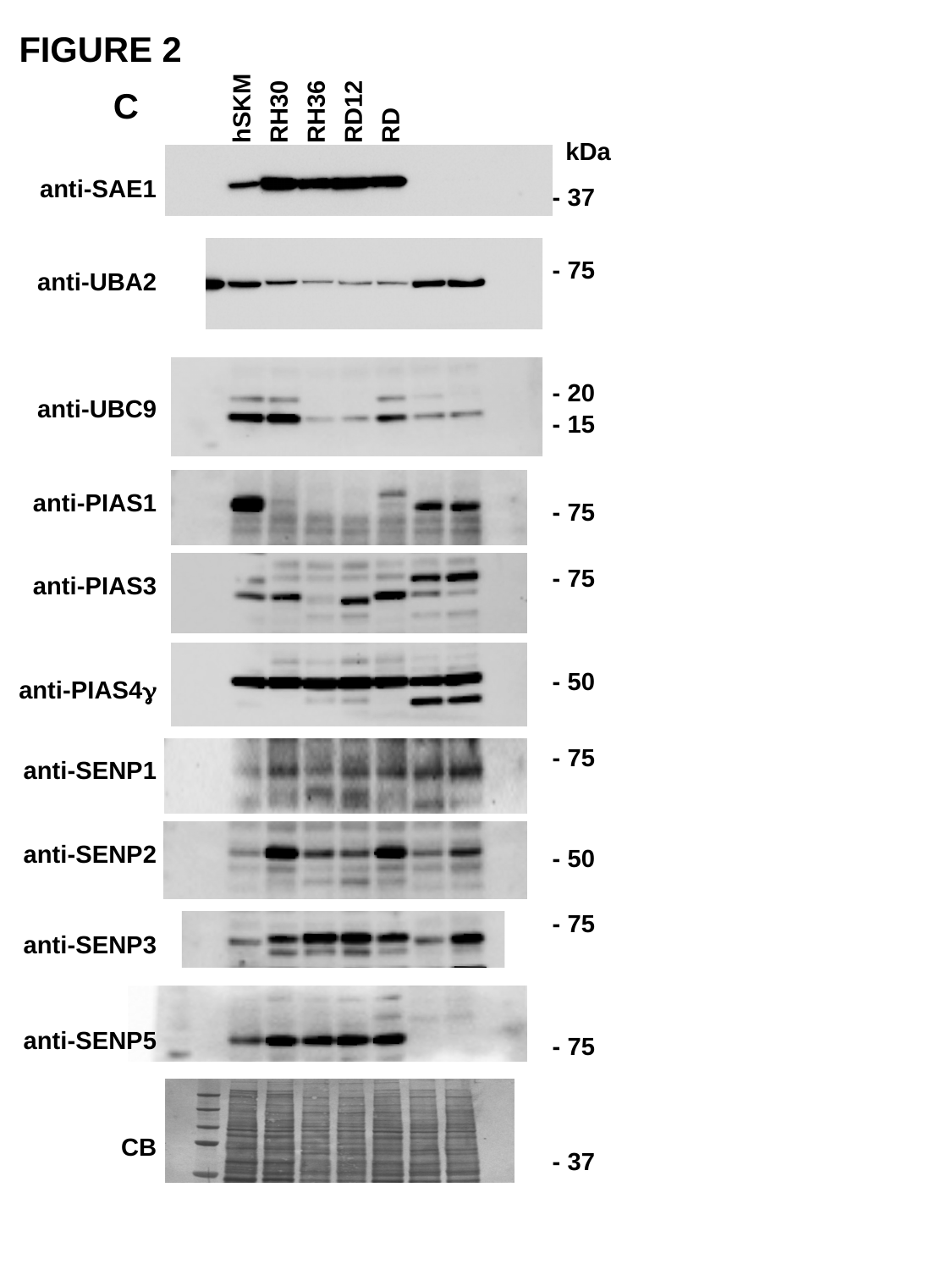

FIGURE 2
RD
hSKM
RD12
RH30
RH36
C
kDa
anti-SAE1
- 37
- 75
anti-UBA2
- 20
anti-UBC9
- 15
anti-PIAS1
- 75
- 75
anti-PIAS3
- 50
anti-PIAS4γ
- 75
anti-SENP1
anti-SENP2
- 50
- 75
anti-SENP3
anti-SENP5
- 75
CB
- 37

### Slide 3
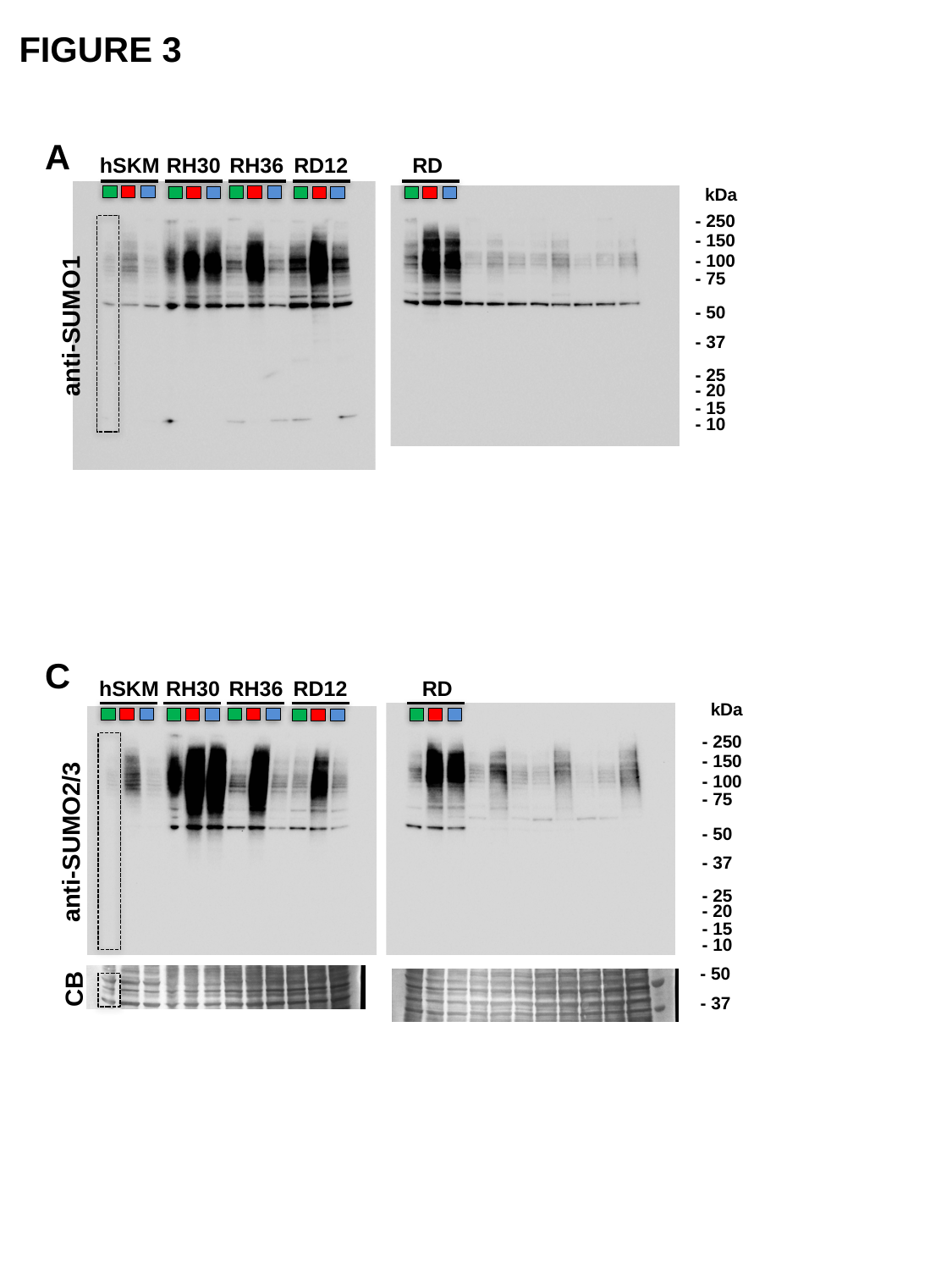

FIGURE 3
A
hSKM
RH30
RH36
RD12
RD
kDa
- 250
- 150
- 100
- 75
- 50
- 37
- 25
- 20
- 15
- 10
anti-SUMO1
C
hSKM
RH30
RH36
RD12
RD
kDa
- 250
- 150
- 100
- 75
- 50
- 37
- 25
- 20
- 15
- 10
anti-SUMO2/3
- 50
CB
- 37

### Slide 4
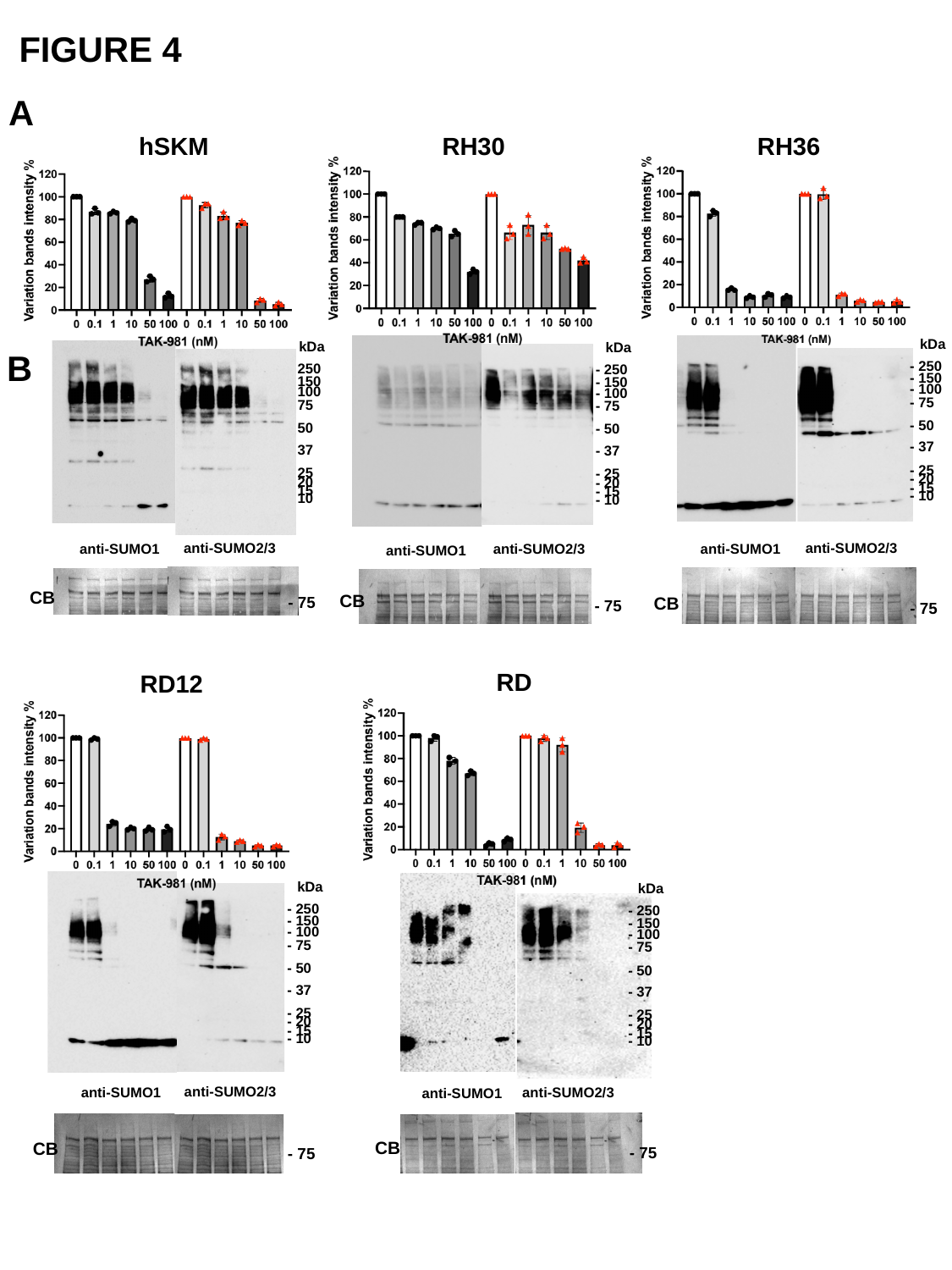

FIGURE 4
A
hSKM
RH30
RH36
kDa
- 250
- 150
- 100
- 75
- 50
- 37
- 25
- 20
- 15
- 10
kDa
- 250
- 150
- 100
- 75
- 50
- 37
- 25
- 20
- 15
- 10
kDa
- 250
- 150
- 100
- 75
- 50
- 37
- 25
- 20
- 15
- 10
B
anti-SUMO2/3
anti-SUMO2/3
anti-SUMO1
anti-SUMO2/3
anti-SUMO1
anti-SUMO1
CB
CB
CB
- 75
- 75
- 75
RD
RD12
kDa
- 250
- 150
- 100
- 75
- 50
- 37
- 25
- 20
- 15
- 10
kDa
- 250
- 150
- 100
- 75
- 50
- 37
- 25
- 20
- 15
- 10
anti-SUMO2/3
anti-SUMO1
anti-SUMO2/3
anti-SUMO1
CB
CB
- 75
- 75

### Slide 5
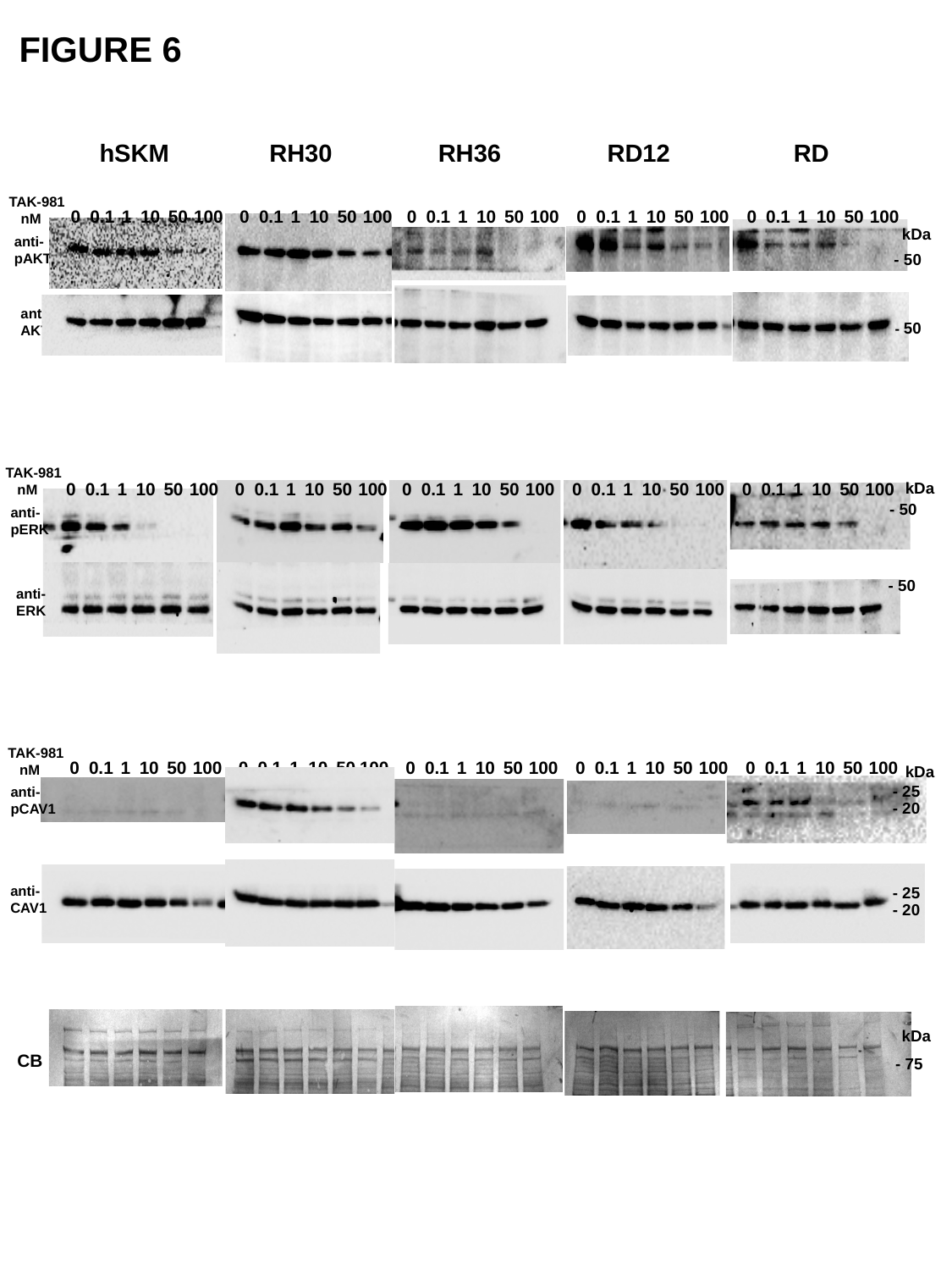

FIGURE 6
hSKM
RH30
RH36
RD12
RD
 TAK-981 nM
0
0.1
1
10
50
100
0
0.1
1
10
50
100
0
0.1
1
10
50
100
0
0.1
1
10
50
100
0
0.1
1
10
50
100
kDa
anti-pAKT
- 50
anti-AKT
- 50
 TAK-981 nM
0
0.1
1
10
50
100
0
0.1
1
10
50
100
0
0.1
1
10
50
100
0
0.1
1
10
50
100
0
0.1
1
10
50
100
kDa
- 50
anti-pERK
- 50
anti-ERK
 TAK-981 nM
0
0.1
1
10
50
100
0
0.1
1
10
50
100
0
0.1
1
10
50
100
0
0.1
1
10
50
100
0
0.1
1
10
50
100
kDa
- 25
anti-pCAV1
- 20
anti-CAV1
- 25
- 20
kDa
CB
- 75

### Slide 6
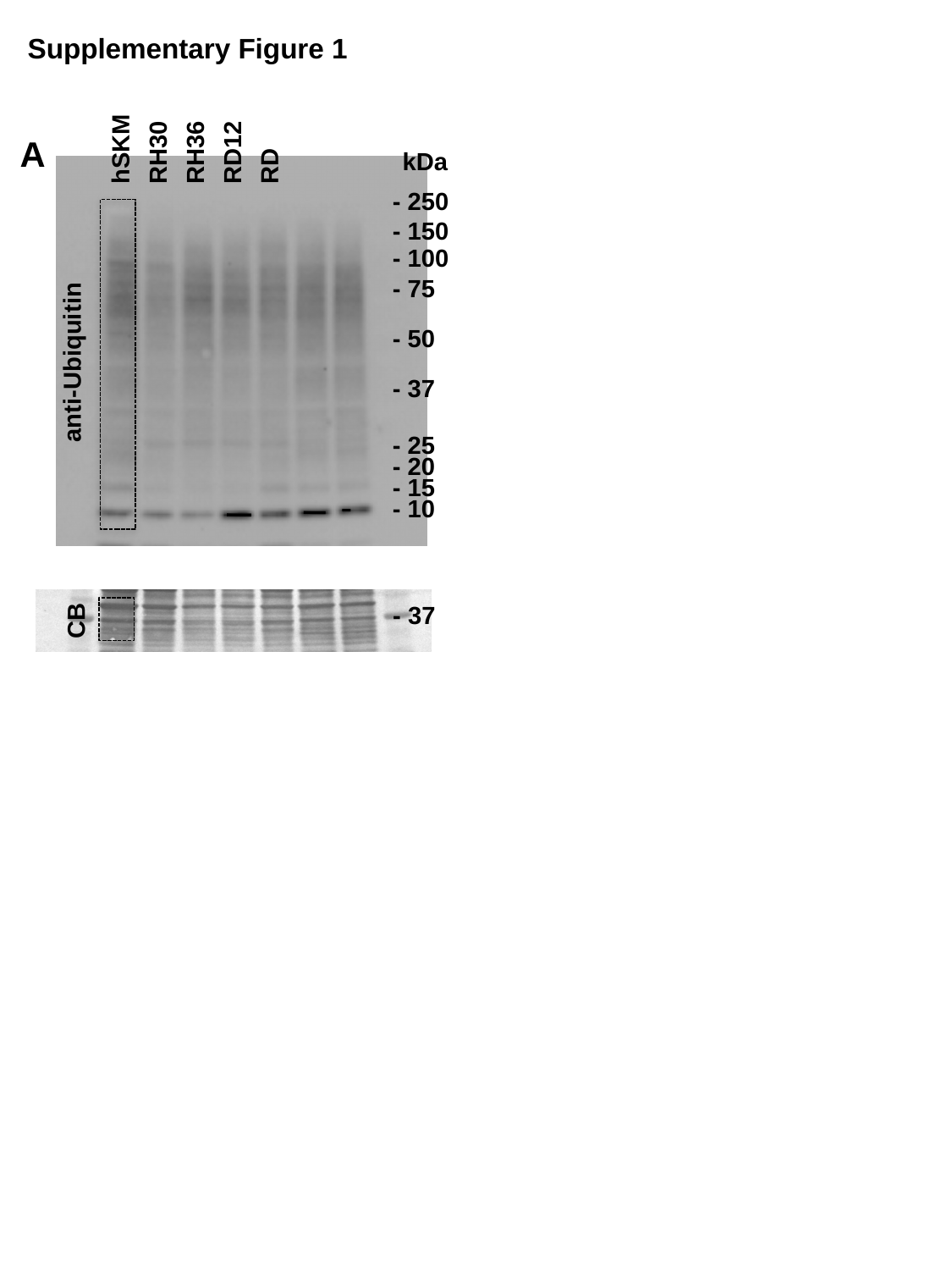

Supplementary Figure 1
RD
hSKM
RH30
RH36
RD12
A
kDa
- 250
- 150
- 100
- 75
- 50
anti-Ubiquitin
- 37
- 25
- 20
- 15
- 10
- 37
CB

### Slide 7
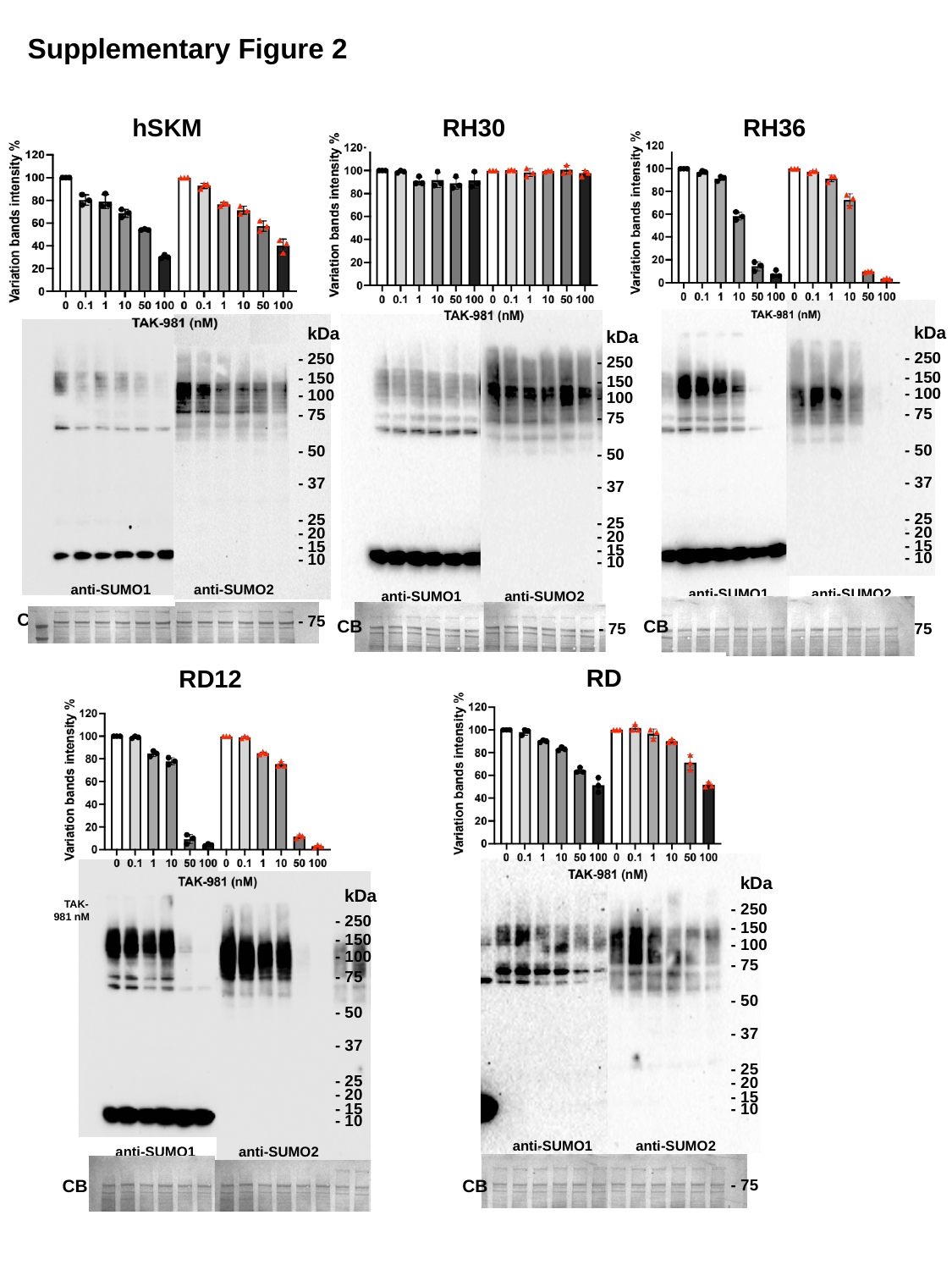

Supplementary Figure 2
hSKM
RH30
RH36
kDa
- 250
- 150
- 100
- 75
- 50
- 37
- 25
- 20
- 15
- 10
kDa
- 250
- 150
- 100
- 75
- 50
- 37
- 25
- 20
- 15
- 10
kDa
- 250
- 150
- 100
- 75
- 50
- 37
- 25
- 20
- 15
- 10
anti-SUMO1
anti-SUMO2
anti-SUMO1
anti-SUMO2
anti-SUMO1
anti-SUMO2
CB
- 75
CB
CB
- 75
- 75
RD
RD12
kDa
- 250
- 150
- 100
- 75
- 50
- 37
- 25
- 20
- 15
- 10
kDa
- 250
- 150
- 100
- 75
- 50
- 37
- 25
- 20
- 15
- 10
 TAK-
981 nM
anti-SUMO1
anti-SUMO2
anti-SUMO1
anti-SUMO2
CB
- 75
CB
- 75
